# Sorting the mob: using geometric morphometrics and machine learning to differentiate kangaroo postcrania for zooarchaeological applications

**DOI:** 10.1101/2024.12.06.627105

**Authors:** Erin Mein, Tiina Manne, Peter Veth, Vera Weisbecker

## Abstract

Taxonomic identification of bone is one of the building blocks of zooarchaeological research into human foraging behaviour. However, it can prove difficult in regions, such as Australia, that have biodiverse taxa that are difficult to differentiate using bone morphology. One such case are the kangaroos and wallabies (macropods), one of the most speciose groups of marsupials whose remains are frequently recovered from Australian archaeological sites. Despite their clear importance to Indigenous economies, little research has been undertaken on how to reliably differentiate the postcranial remains of extant macropods. Here we address this gap by applying three-dimensional geometric morphometrics to describe how the astragalus and calcaneus differs between several large macropod genera. We describe taxonomically diagnostic anatomical attributes between genera and identify the size related (allometric) shape variation that could be mistaken for taxonomic differences. We then compare several machine learning models to demonstrate how these can be applied to geometric morphometric data to statistically classify unknown specimens from palaeozoological contexts with a high degree of accuracy. Our results show that non-linear methods of supervised machine learning outperform classical discriminant function analysis when used on our geometric morphometric data. Statistical classification of palaeozoological specimens has the potential to be a valuable tool, where differentiating skeletal remains of closely related taxa continues to prove challenging.

## Introduction

Applying human behavioural ecology (HBE) models such as prey choice, patch choice and central place foraging to zooarchaeological data requires a detailed understanding of animal behaviour and ecological niche. Kangaroos and wallabies (macropods) were frequently hunted by Indigenous peoples across Australia (Bird et al., 2009; Codding, 2011; Gould, 1969) and the remains of these iconic marsupials are commonly found in archaeological sites (Mein and Manne, 2021). Large macropods (>10 kg) are a particularly interesting group that includes several species that overlap in body size, are wide ranging and sympatric (overlapping in range) but differ in habitat preference, social behaviour and predator escape strategies (Dawson, 2012). High-resolution taxonomic identification (to genus or species) of macropod specimens can therefore provide valuable information on differences in animal ecology and behaviour that influence estimations of handling costs and prey rank (Nagaoka, 2019). At present, high-resolution identification of macropod remains is largely undertaken on craniodental specimens (Mein and Manne, 2021) and comparatively little research has been undertaken on the diagnostic anatomical attributes that differentiate the macropod postcranial skeleton. Limitations on the number of specimens that can be identified beyond the family level has flow on effects for quantifying taxonomic richness (NTAXA), observing patterns in diet breadth and differential body part transport and thus evaluating hypotheses regarding foraging efficiency (Nagaoka, 2019; Grayson, 1989). A better understanding of the anatomical attributes that can be used to differentiate macropod postcrania beyond the family level would therefore advance research into Indigenous foraging behaviour in Australia.

In this study we use geometric morphometrics to describe three-dimensional (3D) shape differences between the astragalus (also referred to as the talus) and calcaneus of several large macropods and evaluate the use of machine learning as a tool for taxonomically identifying morphologically similar specimens. The macropod astragalus and calcaneus are composed of dense, compact bone that, although of low food utility (Lyman, 1992; O’Connell and Marshall, 1989; Garvey, 2010), are known from other mammals to survive well in archaeological contexts (Lyman, 1994).

Prideaux and Warburton (2010) have demonstrated that the macropod ankle reflects evolutionary adaptations for mobility between the upper and lower ankle joints. The ankle morphology of extant large macropod species has been broadly described as reflecting adaptations for reduced mobility for efficient bipedal hopping (Szalay, 1994; Warburton and Prideaux, 2010). Previous descriptive and morphometric studies have largely aimed to infer locomotion in extinct macropods rather than differentiate between the closely related extant taxa (see Bishop, 1997; Flannery, 1982; Janis et al., 2014). However, these studies establish that macropod ankle morphology reflects locomotory adaptations to habitat and substrate that may also exist between extant large macropods.

In a recent study using traditional morphometrics and multiple discriminant analysis, we demonstrated that linear measurements could be used to assign the ankle and foot bones of a wide range of macropods to genus (Mein et al., 2022). Application to macropod bone from Boodie Cave, Barrow Island revealed the co-occurrence of two locally extirpated taxa and adds to the growing body of evidence that the environment of the coastal deserts of northwest Australia was substantially different during the terminal Pleistocene from today. However, in the exact anatomical attributes that differentiate large macropod ankle bones remains unclear. One of the reasons for this was influence of allometry, the covariation of shape with size, that can be related to ontogeny (growth), evolutionary differences between taxa, or intraspecific size differences (e.g. sexual dimorphism) (Klingenberg, 1996, 2016). Although small sample sizes meant we were unable to adequately test for sexual dimorphism in our previous study (Mein et al., 2022), it is likely that age and sexual dimorphism alongside natural intraspecific variation is responsible for the overlap in the linear measurements of large macropod ankle bones. Adult males in the larger macropod genera can be over twice the weight of females (Van Dyck and Strahan, 2008) and foot length in red kangaroos (*Osphranter rufus*), the largest species, has been shown to scale differentially with growth between males and females (Mitchell et al., 2023).

Applying the landmark approach used in geometric morphometrics (GMM) may provide an improvement on the characterisation of taxonomically diagnostic attributes of large macropod ankle bones. GMM uses configurations of landmarks placed on homologous anatomical landmarks within a cartesian coordinate system to describe and compare shape between specimens (Adams et al., 2013). The advantages of using a landmark approach compared to linear measurements is that bone shape is assessed wholistically because spatial relationships between all landmarks are analysed (Weisbecker et al., 2023). Intraspecific variation that is not related to taxonomy, of which allometry can be an important component, is essential to understand when attempting to identify palaeozoological bone (Bochenski, 2008; Driver, 1992). GMM is particularly useful for accounting for allometry as the Procrustes superimposition of landmark configurations allows for the removal of the isometric size – or proportional – component (Goodall, 1991; Viacava et al., 2023). Analyses of allometry can then be undertaken via a regression of shape against size (Klingenberg, 2016). Multivariate analyses of shape differences between *a priori* groups can be applied to geometric morphometric data and importantly for future zooarchaeological applications, three-dimensional (3D) shape differences can be visualised (Klingenberg and Monteiro, 2005).

Machine learning is increasingly being applied as a tool for statistically classifying morphologically similar specimens into taxonomic groups (Hanot et al., 2017; Cole et al., 2022; Moclán et al., 2023; Soda et al., 2017; Salifu et al., 2022). We evaluate the performance of the commonly used linear discriminant analysis (LDA), against three non-linear, supervised machine learning methods: k-Nearest Neighbour (k-NN), Random Forest (RF) and Support Vector Machine (SVM) using our GMM shape data. We discuss the usefulness of machine learning as a tool to support palaeozoological research and as a means of quantifying uncertainties in specimen identification and addressing long-standing challenges of analytical transparency and inter-observer variability in zooarchaeological research.

Taxonomic resolution in mammalian specimen identification is currently a core methodological challenge for Australian zooarchaeology (Mein and Manne, 2021). A narrow focus on craniodental remains and low taxonomic resolution in postcranial specimen identification imposes limitations on our ability to examine patterns in taxonomic richness, patch choice or body part frequency and therefore evaluate hypotheses regarding foraging efficiency and resource depression (Broughton et al., 2010, 2023; Nagaoka, 2019). Our study begins to address these methodological challenges by clarifying where shape differences can be observed between the ankle bones of several large macropod taxa. We provide replicable protocols, using machine learning and GMM data, for statistically classifying those specimens that continue to be hard to visually differentiate. These new data on diagnostic anatomical attributes and GMM and machine learning protocols can be applied to future archaeological specimens to improve the taxonomic identification of large macropod tarsal bones in Australian archaeological assemblages.

## Materials and Methods

We used digital three-dimensional (3D) models of modern astragali and calcanea from 10 species of large macropods as a comparative sample. These species have an average weight of over 10 kg and belong to four genera (Table 1). Unlike evolutionary-focused GMM studies which exclusively use adults, we explicitly aimed to include juvenile, subadult and adult individuals in our sample to best reflect “real world” palaeozoological assemblage composition. Individuals in our sample were categorised into three age groups following Mein et al., (2022). Where possible, both skeletal elements were digitised from the same individual and our sample included the astragalus and calcaneus of 75 individuals. Reference specimens used in our sample are held in the Australian National Wildlife Collection (ANWC), Australian Museum (AM), Museum and Art Gallery of the Northern Territory (MAGNT), Queensland Museum (QM), South Australian Museum (SAM), Western Australian Museum (WAM) and the University of Queensland Zooarchaeology Lab (UQ).

**Table 1.** Sample of modern astragali and calcanea from extant large macropod species.

| **Species** | **No. Astragali** | **No. Calcanea** |
| --- | --- | --- |
| *Macropus fuliginosus* (western grey kangaroo) | 9 | 10 |
| *Macropus giganteus* (eastern grey kangaroo) | 8 | 8 |
| *Notamacropus agilis* (agile wallaby) | 13 | 13 |
| *Notamacropus parryi* (whiptail wallaby) | 1 | 1 |
| *Notamacropus rufogriseus* (red-necked wallaby) | 3 | 3 |
| *Osphranter antilopinus* (antilopine wallaroo) | 6 | 5 |
| *Osphranter bernardus* (black wallaroo) | 5 | 5 |
| *Osphranter robustus* (common wallaroo) | 18 | 18 |
| *Osphranter rufus* (red kangaroo) | 15 | 14 |
| *Wallabia bicolor* (swamp wallaby) | 3 | 3 |
| **Total** | **81** | **80** |

Digital models were produced using a Solutionix C500 or a Shining 3D EinScan Pro+ structured light surface scanner. 3D meshes were produced using the software Ezscan 2017 (Medit, 2019) or EinscanPro v2.6.0.9 (Shining 3D, 2022) respectively and decimated in Meshlab to reduce file size (Cignoni et al., 2008). All 3D meshes are available on the MorphoSource project ‘Sorting the Mob’.

### Landmarking Protocol

We placed a total of 32 fixed landmarks and 21 sliding semi-landmarks on each astragalus and 35 fixed and 13 sliding semi-landmarks on each calcaneus (Supplementary Fig. S1 and Fig. S2). Fixed and sliding semi-landmarks were placed on each 3D mesh by EM using Checkpoint (Stratovan Corporation, 2018). Semi-landmarks were placed equidistantly along curves (Adams et al., 2013) and 3D models of right-sided astragali or calcanea were mirrored in Checkpoint prior to placing landmarks. The calcaneal epiphysis was not included in the landmarking protocol as indeterminate growth patterns in large macropods mean it is frequently missing, even in adult individuals (Dawson, 1995). Additional details of the landmarking protocols can be found in the supplementary information (Supplementary Table S1 and Table S2).

We transformed the raw 3D landmark coordinates using Generalised Procrustes Analysis (GPA) to remove the effects of rotation, translation and size prior to analysis (Rohlf and Slice, 1990). Semi-landmarks were slid along their curves using the minimum bending energy criterion (Bookstein, 1989). GPA rescales coordinate configurations to the mean centroid size (square root of sum of squared distances of all landmarks to the centroid (Bookstein, 1991)). This results in Procrustes shape coordinates which are free of isometric size (the part of size variation where specimens vary by the same amount) and can subsequently be used as shape variables in multivariate analyses.

### Shape differences between groups

We initially explored shape differences between taxonomic groups by performing Principal Components Analysis (PCA) on the Procrustes shape coordinates. We examined the distribution of specimens across the first three principal components (PC) to examine how shape differed between taxa in the main variation of the data. Because PCA is agnostic to group membership, the separation of specimens into taxonomic morphospaces in the main variation would strongly suggest that distinctive differences in shape do occur between taxonomic groups (Strauss, 2010).

### Allometries

We compared the size of each bone by plotting centroid size by species and identified where taxa overlapped in size. We then used Procrustes Analysis of Variance (ANOVA) to test whether bone shape differs significantly with centroid size and taxonomic group. Significant interactions between size and genus indicate differences in allometric slopes which means bone shape differs between groups variably with changing size (Collyer et al., 2015). We performed a pairwise comparison to test where allometric slopes differed significantly between genera and plotted the allometric regression scores against log centroid size to visualise how allometric slopes vary between genera (Collyer et al., 2015; Drake and Klingenberg, 2008). Where we found no significant slope differences (or the interaction term was significant but did not explain much variation compared to the grouping variable), we asked if the genera differed in intercept. Differences in intercept indicates consistent differences in shape are present between groups even where allometric slopes are the same. We were unable to statistically test the relationship between shape and sex due to the absence of this metadata for a large proportion of the sample but expect sex to have relatively little impact on ankle bone shape aside from overall size and allometric patterns.

### Visualisation of shape differences

We visualised shape differences between genera by warping a reference mesh, using the thin-plate spline method (Bookstein, 1989), to the mean shape of each group along the eigenvectors of the first three principal components of a between group PCA (bg-PCA) using genus as the grouping factor. A bg-PCA was used to maximise the separation between the groups, while retaining Procrustes distances in the original shape space (Klingenberg and Monteiro, 2005; Mitteroecker and Bookstein, 2011). We cross-validated all bg-PCAs to avoid issues of spurious group differences (Cardini and Polly, 2020). To visualise allometric variation we warped a reference mesh along the fitted allometric slope of a Procrustes ANOVA on each taxonomic group (Adams and Otarola-Castillo, 2013). We did not correct for allometry to examine only taxonomic differences as taxonomic allometry is an important part of visual comparative identification.

### Statistical Classification

We tested whether bone shape could be used to predict taxonomic group membership to the genus level, using linear and non-linear supervised machine learning models. Shape data from our modern sample was used to ‘train’ each model. The principal components produced in each PCA performed on the Procrustes shape coordinates were used as predictors. This reduces the dimensionality of the dataset and allows for reduction of variables to those that explain the variation sufficiently (see below for details). Statistical classification to species was not attempted due to sample size at the species level. Due to the relatively small size of our training sample, all models were cross-validated using a leave-one-out cross validation repeated 1000 times rather than splitting the sample into training and test populations (Kuhn and Johnson, 2016).

Linear Discriminant Analysis (LDA) is a traditional, linear method of data regression and is widely used for applied classification problems (Kovarovic et al., 2011; Strauss, 2010). However, LDA assumes that the underlying data are normally distributed, that all covariance matrices are equal, and the within-group sample size is higher than the number of predictors (Hair et al., 2018; Mitteroecker and Bookstein, 2011). These assumptions are often violated by real-world biological data and particularly the high-dimensional data of geometric morphometrics (Dryden and Mardia, 1993). LDA models produced with uneven class sizes may result in a high overall classification accuracy but accuracy within each class may be unbalanced (Courtenay, 2023; Sanchez, 1974). Classes with larger populations may achieve high classification accuracy but, accuracy in smaller classes will be lower (Soda et al., 2017). We equalised the prior probabilities across the four classes (genera) (0.25) to reduce the effect of class sample size on the posterior probabilities.

K-Nearest Neighbour (k-NN) algorithms are a simple but effective machine learning method for statistical classification. A ‘lazy learner’ method, k-NN algorithms do not model the training data but rather calculate the distance between unknown specimens and *k* number of nearest neighbours in a training sample (Salifu et al., 2022). A prediction of class membership is made based on the proportion of neighbours in each class in the training sample. Each k-NN algorithm was tuned on *k* between 2 and 20 (Kuhn and Johnson, 2016).

Random Forest (RF) is an ensemble model in which the results of many randomised classification and regression trees (CART) are aggregated to produce a final class prediction (Arriaza and Domínguez-Rodrigo, 2016). The tuning parameter *mtry* sets the number of predictors considered at each decision node and the predictors are randomised in each iteration. Due to their logic-based branch and node structure, RF models make no assumptions about the underlying structure of the data and provide a powerful statistical classification method for high-dimensional and non-normally distributed data (Cole et al., 2022). Following Kuhn and Johnson (2016) the number of individual trees in each RF was set to 1000 and *mtry* was tuned on values between 2 and *p*-1 where *p* is the number of principal components (see below).

Support Vector Machine (SVM) algorithms aim to establish a hyperplane classification boundary between data classes in multidimensional space (Courtenay et al., 2019). We tested a SVM algorithm with a simple linear kernel where sigma (σ), a parameter which smooths the hyperplane, was held constant. A cost (C) parameter is applied which imposes a penalty value on the model for misclassification. Low C values produce a smooth hyperplane which tend to underfit the data, while high C values will produce a hyperplane which is overfitted to the training data and cannot be generalised to new data (Kuhn and Johnson, 2016). We tuned the algorithm on a C parameter between 2^±20^. SVM algorithms are a powerful and flexible method of machine learning for statistical classification which have been demonstrated to work well with the highly dimensional data that is common to geometric morphometrics (Salifu et al., 2022).

As each of these machine learning models can be sensitive to uninformative predictors, principal components were iteratively added to the models and the number of principal components which optimised Accuracy and Kappa scores were included in the final models (Baylac and Frieß, 2005; Kuhn and Johnson, 2016). Reduction of model accuracy beyond the optimised number of principal components indicates that shape information held in additional components is unlikely to relate to taxonomic differences between groups and may introduce uninformative ‘noise’ to the models. The number of principal components iteratively added to the LDA models were limited to the first 16 to ensure the number of predictors did not exceed the within-group sample size (except *Wallabia*) (Mitteroecker and Bookstein, 2011).

We compared the overall accuracy scores and rate of misclassification within each taxonomic group (within-group sensitivity) for each model. The adequacy of each model for application to real world palaeozoological data was assessed against the 1.25 x maximum chance (Cmax) criterion proposed by Hair et al., (2018). If no information is provided, the best chance at correctly classifying the maximum number observations is to classify all observations to the largest group. Therefore, the Cmax criterion is the proportion of the largest group in the overall sample.

All analyses were undertaken in R v.4.2.2 (R Development Core Team, 2023) using the packages *geomorph* (Adams et al., 2022), *Morpho* (Schlager et al., 2021), *caret* (Kuhn, 2022) and *landvR* (Guillerme and Weisbecker, 2019).

## Results

### Astragali

#### Principal Components Analysis

Distribution of specimens in the main variation of the PCA (PC1-PC2) on astragalus shape reveal very little overlap between the morphospace of *Macropus*, *Notamacropus* and *Osphranter.* This indicates that the astragali of these three genera are distinct from each other, but *Wallabia* astragali may be very similar in shape to *Notamacropus* (Fig. 1). Juvenile specimens in the *Osphranter* and *Macropus* genera tended to score highly along PC2 which indicates some ontogenetic allometry may be observed in the astragali of these two genera.

**Fig. 1.**
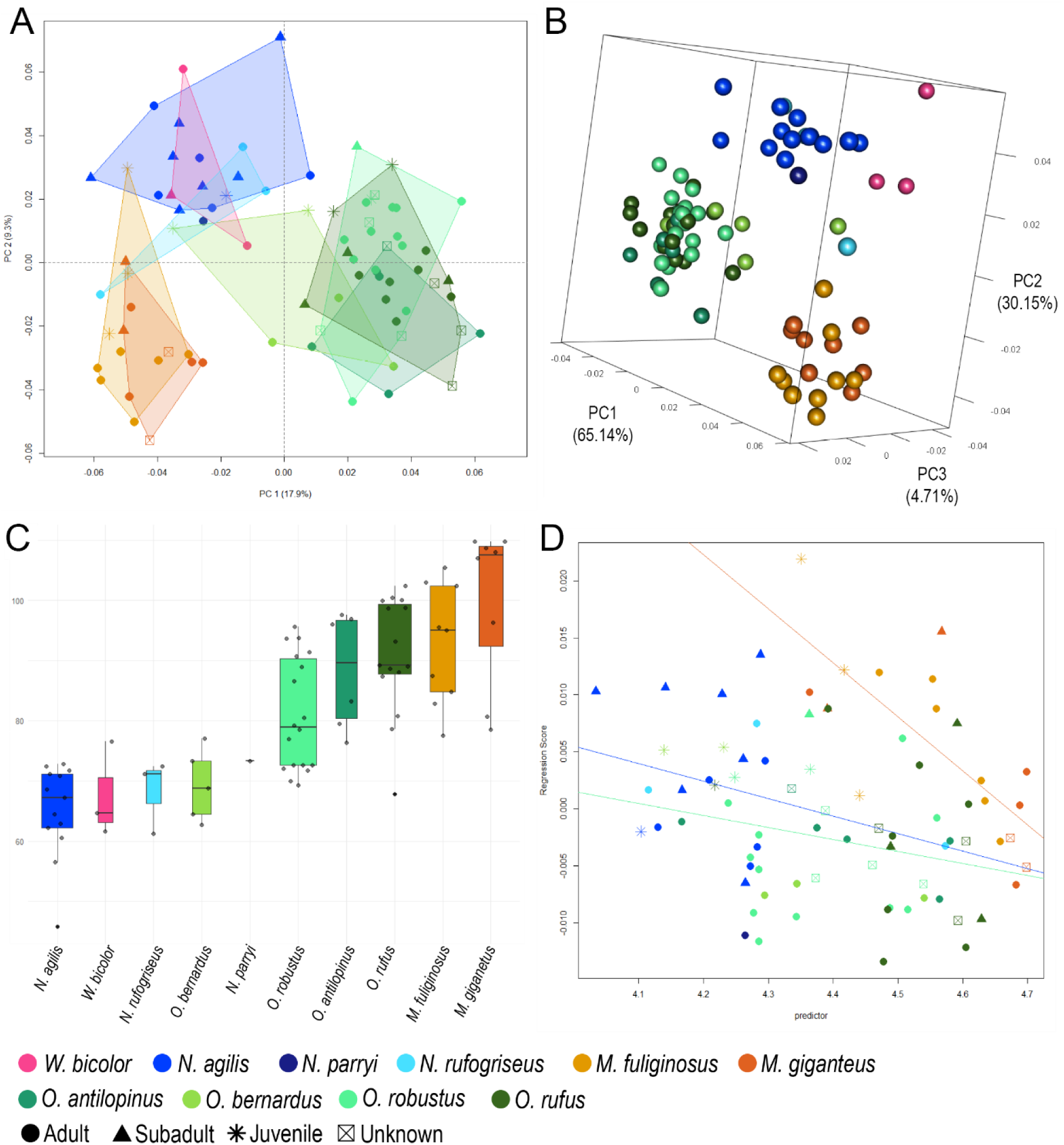
Results of analyses on large macropod astragali. **a)** Main variation of the PCA (PC1-PC2, 27.2% variance) **b)** Main variation of the bg-PCA (PC1-PC3, 100% variance); **c)** comparative centroid size of astragali by species; and **d)** Plot of the allometric regression score against log(centroid size).

Macropods that occupy the most open environments, such as *O. rufus* (arid grasslands) (Dawson, 2012) and *O. antilopinus* (tropical savannah) (Ritchie, 2007), occupy the positive PC1, negative PC2 shape space (Fig. 1). Conversely *Notamacropus* and *W. bicolor* which live in the most closed environments (forests) occupy the opposing negative PC1, positive PC2 shape space and the *Macropus* specimens (open woodland) fall between these two extremes (Van Dyck and Strahan, 2008). The morphospace of *O. bernardus,* which inhabits rocky escarpments (Press, 1989), falls closer to the *Notamacropus/Wallabia* morphospace along PC3. Although specimens are grouped strongly by taxonomy in the main variation of the PCA, the distribution of groups across the first three principal components suggests that astragalus shape may also reflect habitat structure.

#### Procrustes ANOVA

The Procrustes ANOVA indicates that size accounts for comparatively little variation in astragalus shape (6.3%) in our sample compared to the differences accounted for by taxonomy (20.2%) (Table 2). The interaction term between size and genus was not significant indicating the shape of *Macropus*, *Notamacropus* and *Osphranter* astragali follow a similar allometric slope (Supplementary Table S3). Differences in intercept indicates that despite astragali shape varying with size consistently amongst the *Macropus*, *Notamacropus* and *Osphranter*, consistent differences in shape are still present between the astragali of these three genera at all sizes (Fig. 1).

**Table 2.** Results of the Procrustes ANOVA performed on ankle bone shape against centroid size and genus.

|  | Df | SS | MS | Rsq | F | Z | Pr(>F) |
| --- | --- | --- | --- | --- | --- | --- | --- |
| *Astragalus* |  |  |  |  |  |  |  |
| Shape ~ Size | 1 | 0.035226 | 0.035226 | 0.062874 | 6.292409 | 4.576023 | 0.001 |
| Shape ~ Genus | 2 | 0.11332 | 0.05666 | 0.202262 | 10.12112 | 6.63952 | 0.001 |
| Shape ~ Size * Genus | 2 | 0.014246 | 0.007123 | 0.025428 | 1.272408 | 1.55874 | 0.067 |
| Residuals | 71 | 0.397471 | 0.005598 | 0.709436 | NA | NA | NA |
| *Calcaneus* |  |  |  |  |  |  |  |
| Shape ~ Size | 1 | 0.085421 | 0.085421 | 0.1343 | 14.68758 | 4.829349 | 0.001 |
| Shape ~ Genus | 2 | 0.116867 | 0.058433 | 0.183738 | 10.04716 | 5.089433 | 0.001 |
| Shape ~ Size * Genus | 2 | 0.020834 | 0.010417 | 0.032755 | 1.791096 | 2.944415 | 0.003 |
| Residuals | 71 | 0.412929 | 0.005816 | 0.649208 | NA | NA | NA |

#### Visualising Shape Differences

The three-dimensional visualisation of shape differences between genera in the bgPCA reveals key differences in astragalus shape between the large macropods (Fig. 2). *Macropus* specimens have broad (medio-lateral) and short (antero-posterior) trochlear crests. When viewed superiorly, these give the astragalus a square appearance. The medial trochlear crest in *Macropus* specimens is positioned more posterior to the navicular facet and the malleolar fossa is comparatively small and circular. The other three genera have more obliquely orientated trochlear crests. *Osphranter* astragali are distinguished by their long medial plantar tuberosity and narrow, tapered trochlear crests which give the astragalus a triangular appearance in plan. The curvature of the lateral talocalcaneal facet is also comparatively shallow on *Osphranter* astragali. *Notamacropus* and *Wallabia* astragali have broad but obliquely orientated trochlear crests, the medial crest is positioned lateral to the navicular facet and the lateral crest projects more anteriorly than in *Macropus*. The medial malleolar fossa is comparatively large, oval in plan, and positioned more anteriorly in *Notamacropus* and *Wallabia*.

**Fig. 2.**
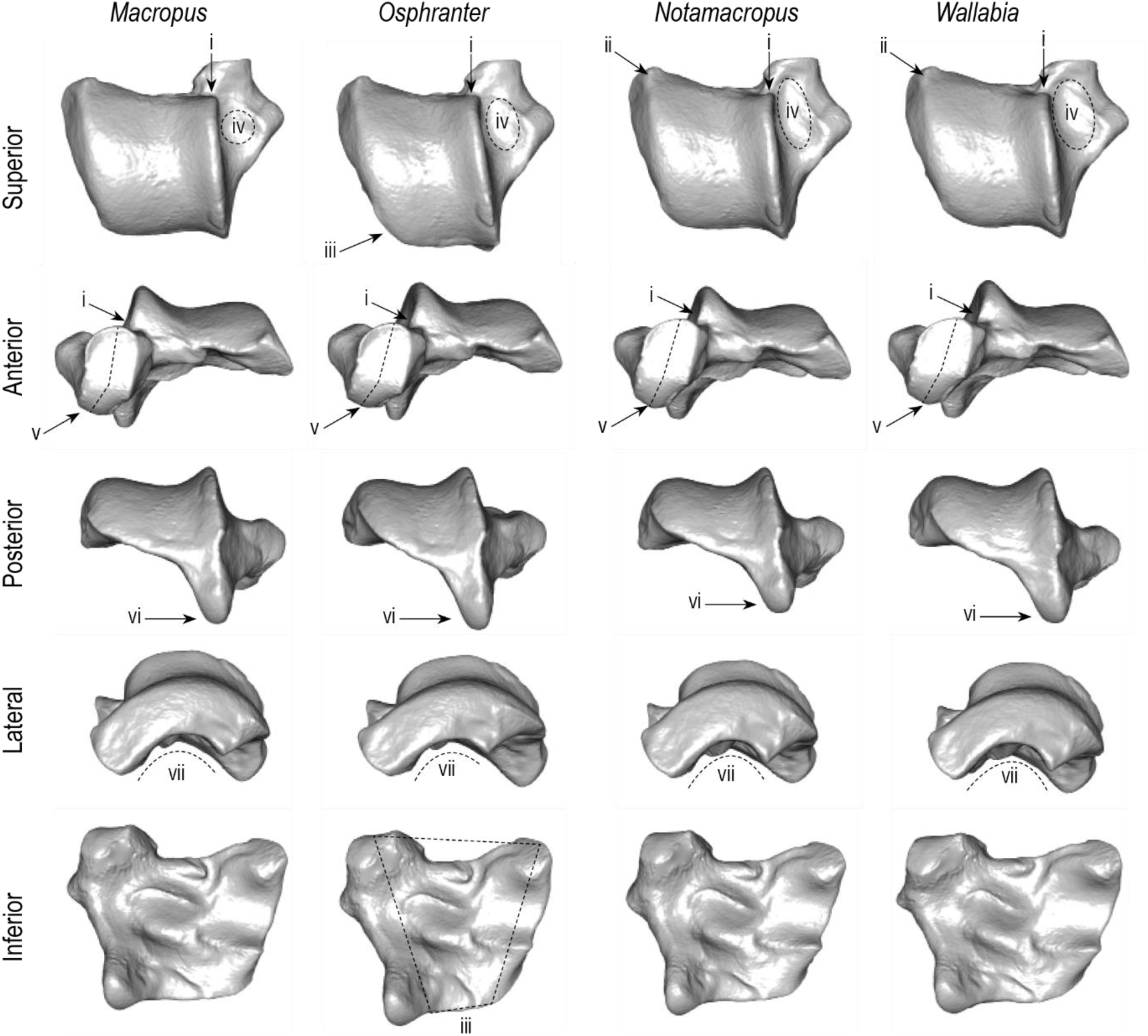
Taxonomically diagnostic characteristics between large macropod astragali. Shape differences observed by warping a reference mesh between mean genus shapes in a cross validated bgPCA using the thin-plate spline method. **i)** medial trochlear crest positioned more posterior to astragalar head in *Macropus*; **ii)** lateral trochlear crest projects more anteriorly in *Notamacropus* and *Wallabia*; **iii)** trochlear crests are elongated and tapered in *Osphranter* creating a triangular appearance in plan; **iv)** medial malleolar fossa is small and circular in *Macropus* and elongated and oval in *Notamacropus* and *Wallabia*; **v)** plantar half of the navicular facet is orientated more medially in *Macropus* and *Osphranter*; **vi)** medial plantar tuberosity is elongated in *Osphranter* and shortest in *Notamacropus* and *Wallabia*; and **vii)** the curvature of lateral talocalcaneal facet is comparatively shallow in *Osphranter*.

Visualisation of allometric variation within each genus shows that the navicular articular facet and talocuboid and secondary talocalcaneal facet on the lateral surface of the astragalar head become taller and broader as size increases (Supplementary Fig. S3 and Video S3). The medial malleolar fossa becomes deeper, and the medial plantar tuberosity is elongated with increasing size, particularly in *Osphranter* and *Macropus* taxa. The curvature of the medial trochlear crest projects more anteriorly and the lateral trochlear crest increases in height in larger *Osphranter* and *Macropus* astragali. On the plantar surface of the astragalus both talocalcaneal articular facets are more open, and less acutely concave in smaller individuals.

#### Statistical Classification

The overall accuracy scores for all models were very high (>0.88) and exceeded the Cmax threshold proposed by Hair et al., (2018) for minimum model adequacy by at least 21% (Table 3). Kappa scores for all four models were over 0.8 indicating very strong agreement between the predicted and observed classes (Kuhn and Johnson, 2016). The k-NN and SVM algorithms produced identical overall model accuracies, but the within-group sensitivity was more balanced in the SVM model meaning the rates of misclassification were more equal between genera. *Macropus* and *Osphranter* astragali were consistently classified correctly across all four methods but in the LDA, k-NN and RF models this was largely achieved via misclassification of *Notamacropus* astragali. The LDA model had the highest rate of misclassification of *Notamacropus* specimens. Owing to the small sample size, *Wallabia* astragali could not be correctly classified to genus in any of the models evaluated here and these specimens were consistently misclassified as *Notamacropus* (Supplementary Table S4). Except for the k-NN algorithm, the inclusion of principal components beyond the first seven to 10 (54.3% to 62.9% variance) resulted in a decline in model performance (Supplementary Fig. S5). This indicates that shape information held in these additional principal components is largely taxonomically uninformative and adds unnecessary noise to the models.

**Table 3.** Overall model accuracy and within-group sensitivity scores by skeletal element and machine learning model.

| Model Sensitivity | LDA | k-NN | RF | SVM |
| --- | --- | --- | --- | --- |
| *Astragali* |  |  |  |  |
| *Macropus* | 1.00 | 1.00 | 0.941 | 1.00 |
| *Notamacropus* | 0.706 | 0.824 | 0.882 | 1.00 |
| *Osphranter* | 0.977 | 1.00 | 0.977 | 0.932 |
| *Wallabia* | 0.000 | 0.000 | 0.000 | 0.000 |
| **Overall Model Accuracy** | **0.889** | **0.926** | **0.914** | **0.926** |
| Standard Deviation | 0.164 | 0.102 | 0.048 | 0.039 |
| Kappa | 0.823 | 0.877 | 0.859 | 0.881 |
| Cmax threshold x 1.25 | 0.679 | 0.679 | 0.679 | 0.679 |
| Optimised No. Predictors | 7 | 21 | 10 | 7 |
| % shape variance included | 54.28% | 81.83% | 62.85% | 54.28% |
| Optimised Tuning Variable | - | 11<sup>a</sup> | 3<sup>b</sup> | 3.27 x 10<sup>-4 c</sup> |
| *Calcaneus* |  |  |  |  |
| *Macropus* | 1.00 | 1.00 | 0.889 | 1.00 |
| *Notamacropus* | 0.765 | 0.941 | 0.941 | 0.882 |
| *Osphranter* | 1.00 | 0.952 | 0.976 | 0.976 |
| *Wallabia* | 1.00 | 0.00 | 0.00 | 0.00 |
| **Overall Model Accuracy** | **0.950** | **0.925** | **0.913** | **0.925** |
| Standard Deviation | 0.118 | 0.031 | 0.044 | 0.062 |
| Kappa | 0.921 | 0.881 | 0.859 | 0.879 |
| Cmax threshold x 1.25 | 0.656 | 0.656 | 0.656 | 0.656 |
| Optimised No. Predictors | 9 | 6 | 5 | 5 |
| % shape variance included | 67.59% | 59.11% | 55.32% | 55.32% |
| Optimised Tuning Variable | - | 5<sup>a</sup> | 2<sup>b</sup> | 2.07431<sup>c</sup> |

<sup>a</sup> *k* neighbours<sup>b</sup> *mtry*<sup>c</sup> Cost

### Calcanea

#### Principal Components Analysis

Very little overlap is observed between the morphospace of genera in the main variation of the PCA (PC1-PC2) on calcaneus shape indicating that this bone may also be taxonomically distinct in shape (Fig. 3). Notably, the calcanea of *O. rufus* form an entirely separate morphospace across the first three principal components, suggesting the calcanea of this species may be distinct in shape from the other *Osphranter* species. Again, the *Wallabia* calcanea overlap entirely with the *Notamacropus* morphospace. Juvenile and subadult specimens score more highly along PC1 indicating some ontogenetic allometry may be observed in calcanea of all genera.

**Fig. 3.**
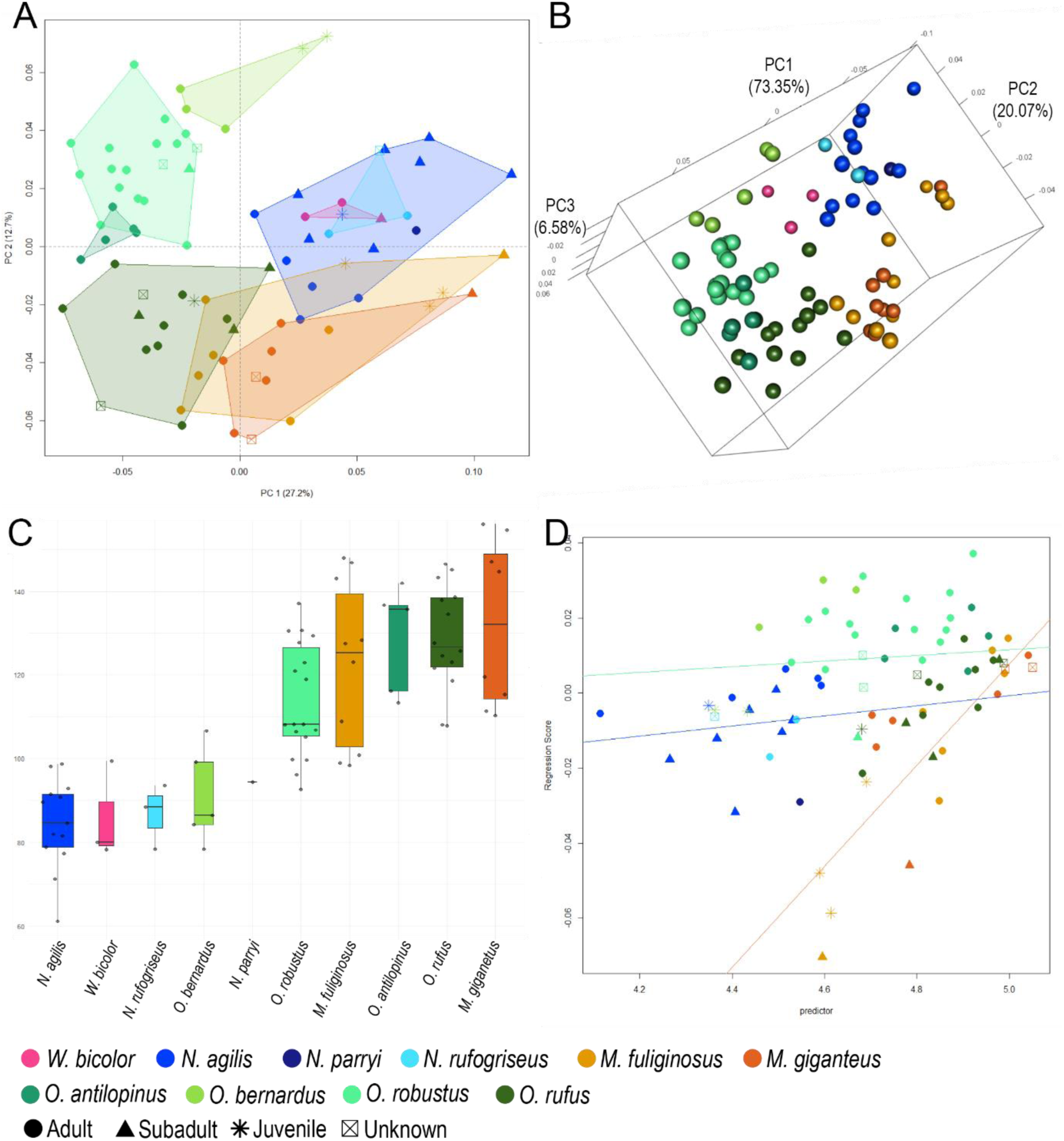
Results of analyses on large macropod calcanea. **A)** Main variation of the PCA (PC1-PC2, 39.9% variance) **B)** Main variation of the bg-PCA (PC1-PC3, 100% variance); **C)** comparative centroid size of calcanea by species; and **D)** Plot of the allometric regression score against log(centroid size).

The distribution of groups in the main variation of the PCA, particularly in the *Osphranter* genus, suggests that the relationship between bone shape and habitat structure may be stronger in the calcaneus than the astragalus. Taxa which typically live in the most open environments (*Macropus* and *O. rufus*), occupy the negative PC2 shape space and those which occupy the most complex, heterogeneous environments (*Notamacropus* and *Wallabia*) fall within the positive PC2 shape space (Fig. 3) (Dawson, 2012; Van Dyck and Strahan, 2008). The *Osphranter* species fall in a spectrum from most open to most complex environments along the PC2 eigenvector with *O. bernardus* (rocky escarpments) and *O. robustus* (hillslopes) (Press, 1989) falling closest to the forest dwelling *Notamacropus and Wallabia* calcanea.

#### Procrustes ANOVA

Results of the Procrustes ANOVA indicates that size accounts for more shape variation in large macropod calcanea (13.4%) than it does in the astragali, although taxonomy continues to be the stronger predictor of shape (18.4%) (Table 2). A significant interaction term between size and genus was observed and a pairwise comparison indicates that *Osphranter* calcanea may have a significantly different allometric trajectory to both *Notamacropus* and *Macropus* (Supplementary Table S3). However, a plot of the allometric regression scores against log centroid size reveals that *O. rufus* calcanea follow a similar allometric slope to both *Macropus* species, while the other *Osphranter* species follow a similar slope to the *Notamacropus*, albeit with differences in intercept (Fig. 3). The slight convergence in allometric slopes with increasing size suggests that the very largest *Osphranter* and *Macropus* specimens exhibit little allometric differences.

#### Visualising Shape Differences

Because the morphospace of *O. rufus* continued to be distinct from the other *Osphranter* species in the cross validated bgPCA and the evidence for a different allometric pattern in this species, we separated *O. rufus* specimens as their own group in the shape visualisations. The tuber calcanei was observed to be more elongated across all *Osphranter* species but it is slightly taller (dorso-plantarly) in *O. rufus* (Fig. 4). *Notamacropus* have the shortest tuber calcanei and the sustentaculum tali projects more medially in this genus and the *Wallabia*, and medio-plantarly in *Macropus* specimens. When viewed medially, the sustentaculum tali is elongated (antero-posteriorly) and shallow (dorso-plantarly) in *O. rufus* and this characteristic is even more pronounced in the other *Osphranter* species. The curvature of the lateral calcaneotalar articular facet is longer and flatter in all *Osphranter* taxa. The dorsolateral calcaneocuboid facet is slightly plantarly deflected in *Macropus*, *Notamacropus* and *Wallabia* calcanea.

**Fig. 4.**
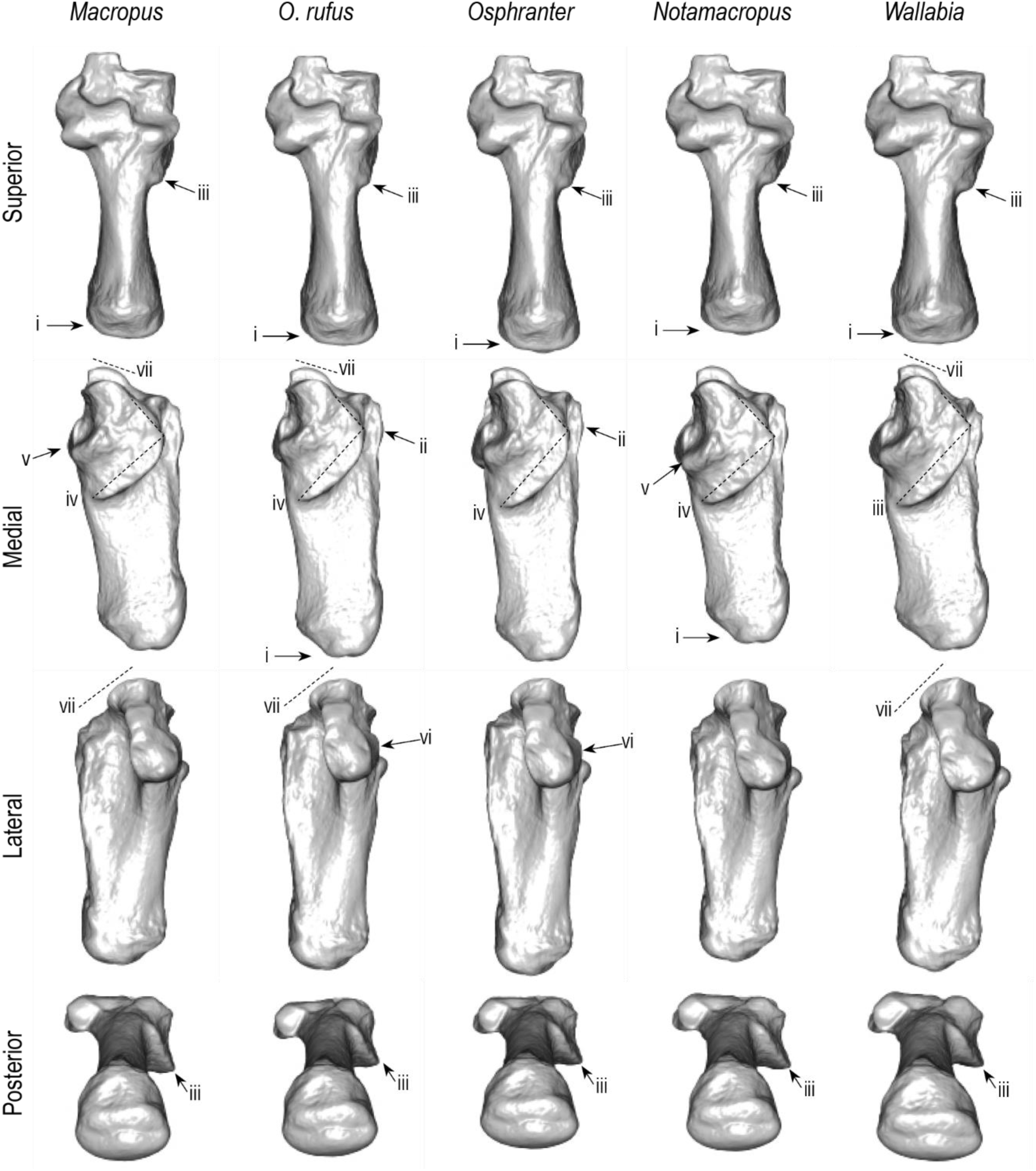
Taxonomically diagnostic characteristics between large macropod calcanea. Shape differences observed by warping a reference mesh between mean genus shapes in a cross validated bgPCA using the thin-plate spline method. **i)** tuber calcanei is elongated in *Osphranter* and shortest and most robust in *Notamacropus* and *Wallabia*; **ii)** tuber calcanei is taller (dorso-plantarly) in *O. rufus* compared to other *Osphranter* species; **iii)** sustentaculum tali projects medially in *Notamacropus* and *Wallabia* and medio-plantarly in *Macropus* and is narrowest in the *Osphranter* genus; **iv)** sustentaculum tali is elongated (antero-posteriorly) and shallow (dorso-plantarly) in the *Osphranter* genus although this is slightly less pronounced in *O. rufus*; **v)** medial calcaneotalar facet projects more dorsally in *Macropus* and is angled more posteriorly in *Notamacropus*; **vi)** curvature of the lateral calcaneotalar facet is longer and flattened in the *Osphranter*; and **vii)** dorsolateral calcaneocuboid facet is deflected plantarly in *Macropus*, *Notamacropus* and *Wallabia*.

Visualisation of allometric patterns shows that generally, small individuals in all genera have a proportionally shorter tuber calcanei and proportionally wider calcaneocuboid and calcaneotalar articulations (Supplementary Fig. S4 and Video S4). In particular, the medial calcaneotalar articular facet is wider and projects more medially in smaller individuals. As indicated by the pairwise comparisons of allometric slopes, small variations to this allometric pattern do occur between the genera but these broad patterns are visually obvious.

#### Statistical Classification

As found in our statistical classification of the astragali, overall accuracy scores for each model in the statistical classification of the calcaneus were very high, and all exceed the Cmax threshold by at least 26% (Table 3) (Hair et al., 2018). As with the astragalus, Kappa scores for all four models were again over 0.8 indicating very strong agreeance between predicted and observed classes (Kuhn and Johnson, 2016). The k-NN and SVM algorithms again produced an identical overall classification accuracy, but the k-NN produced slightly more balanced within-group classification meaning misclassification rates were more equal between genera. The LDA model produced a very high overall classification accuracy and correctly classified all *Macropus*, *Osphranter* and *Wallabia* calcanea but, as with the astragalus, performed worst at correctly classifying *Notamacropus* calcanea. *Wallabia* calcanea were consistently misclassified as *Notamacropus* owing to the small sample and shape similarities (Supplementary Table S5). The inclusion of principal components beyond the first five to nine (55.3% to 67.6% variance) resulted in a decline in model performance. This indicates that shape information held in these additional principal components is largely taxonomically uninformative (Supplementary Fig. S6) and adds uninformative noise to the models.

## Discussion

Low-resolution (family or above) taxonomic identification of terrestrial prey types in Australia currently imposes limitations on our ability to evaluate proposed hypotheses of human foraging behaviour (see O’Connell and Allen, 2012) using zooarchaeological data. Our goal in undertaking this study was to begin to address these methodological gaps by better characterising taxonomically diagnostic attributes that differentiate postcranial bones of common macropods. Comparison of anatomical attributes to infer the taxonomic identity of palaeozoological specimens continues to be a necessity in many regions of Australia as environmental conditions can rapidly destroy the collagen necessary for proteomic or DNA identification (Aplin et al., 2016; Peters et al., 2021; 2023). Most marsupial postcranial remains are currently identified to the family level or higher, providing little data on potential behavioural or ecological differences between prey items. In regions with relatively low biodiversity, identification of specimens to the family level may provide sufficient data to infer prey rank or patch choice. However, in biodiverse regions like Australia, morphologically similar taxa can occupy different ecological niches and exhibit differences in their social, feeding and predator avoidance behaviour. These differences can have substantial flow on effects for recognising patch choice and estimating handling costs and energy returns.

The absence of truly large prey (>100 kg) in Australia (except for extinct megafauna) may require a reconsideration of body size as a correlate for prey rank. Broughton et al. (2011) have suggested that relatively small differences in body size between prey items requires a greater consideration of factors that impact handling costs such as speed, defence behaviours or the capacity for mass harvesting (see also Morin et al., 2019; Stiner et al., 2000). Macropods pose just such a challenge for research into Indigenous foraging behaviours in Australia. For example, grey and red kangaroos (*Macropus* spp.), antilopine wallaroos *(O. antilopinus)* and agile wallabies (*N. agilis*) typically live in gregarious social groups and are diurnal or crepuscular feeders (Van Dyke and Strahan, 2008; Ritchie, 2007). Red kangaroos are more arid adapted and typically occupy flatter lowlands while common wallaroos (*O. robustus*) and black wallaroos (*O. bernardus*) prefer hilly country or rocky escarpments and are typically more solitary. The *Macropus*, *Notamacropus* and *Wallabia* species in our sample prefer woodlands, forests or riverine corridors, although differ in their feeding strategies and diurnal activity levels (Van Dyke and Strahan, 2008). Large kangaroos and wallaroos have the capacity to escape predators at high speeds between 40 to 65 km per hour (Freedman et al., 2020; Hopwood and Butterfield, 1990; Dawson and Taylor, 1973) while the smaller wallaby taxa tend to have slower hopping speeds of approximately 13 km per hour (Bennett, 1987; Baudinette, 1977). These differences in habitat preference, behaviour and speed may be important considerations for inferring prey rank but require specimens to be identified beyond the family level before any patterns can be observed in the zooarchaeological data.

Following the extinction of the megafauna, large macropods are among the very few non-volant terrestrial prey types in Australia to weigh over 10kg (along with wombats and emus). Large macropods likely required field processing prior to transport (O’Connell and Marshall,1989; Bird et al., 2009) and therefore present one of the few opportunities in this ‘continent of small game’ (Bird et al. 2013) to observe differential body part transport in zooarchaeological data (Nagaoka, 2019). Large macropod remains are therefore important for understanding foraging efficiency as well as prey and patch choice in Australia. Yet, skeletal element frequencies are rarely reported in the Australian literature (Mein and Manne, 2021), possibly owing to the current reliance on craniodental specimens for more specific taxonomic identification. Development of methods that allow postcranial specimens of large macropods to be identified beyond the family level increases overall sample sizes of identified specimens and creates opportunities to examine foraging efficiency in relation to evidence for palaeoenvironmental change or resource depression.

Application of GMM in this study has allowed us to summarise and describe the anatomical attributes that taxonomically differentiate large macropod astragali and calcanea beyond what is possible using only linear measurements. Taxonomic identification of palaeozoological specimens requires a strong understanding of non-taxonomic morphological variation, of which allometry is particularly important because it is difficult to identify (Bochenski, 2008; Viacava et al., 2023). Although we did not attempt to separate the causes of allometry (ontogeny, sex, evolution) the visualisation of allometric variation here has allowed us to identify important anatomical attributes that vary with size on both skeletal elements.

We found that the shape of large macropod astragali varied less with size than the calcanea, despite most elements being sourced from the same individuals and a slightly higher proportion of juveniles in the astragalus sample. The forces imposed on the tuber calcanei by the gastrocnemius muscle during hopping are likely responsible for the increased influence of size on calcaneus shape compared to the astragalus. Although allometry has a relatively small effect on total shape variation for both skeletal elements, analysts should consider the allometric characteristics described here when attempting to visually compare the anatomical attributes of individual specimens. For example, the comparative length of the medial plantar process on the astragalus and tuber calcanei on the calcaneus are taxonomically diagnostic characteristics between genera but can also vary with size. This is particularly important where specimens fall within the range of size overlap between juvenile large taxa and adult smaller taxa.

Because of the potential interaction between taxonomic and allometric shape variation, individual anatomical attributes should not be used in isolation but rather in conjunction with attributes from across the entire bone. This may limit our ability to identify fragmented astragali and calcanea beyond the family level where few anatomical attributes are preserved and requires further testing. As with all analytical protocols, further quality assessment of the diagnostic anatomical attributes identified here should be undertaken (Wolverton, 2013).

Blind tests should be conducted to establish the inter-operator reliability of the morphological characteristics identified in this study (see Morin et al., 2017; Lubinski et al., 2020; Zeder and Lapham, 2010).

Our study corroborates previous research that macropod ankle morphology reflects locomotor adaptations to habitat structure (Prideaux and Warburton, 2010). Characteristic features of adaptation to open habitats have been described as a narrow and transversely orientated lower ankle joint between the astragalus and calcaneus, increased area of contact between the lateral astragalar head with the cuboid and calcaneus, an elongated tuber calcanei, a narrow sustentaculum tali and a steeply stepped calcaneocuboid morphology (Barnett, 1970; Bishop, 1997; Janis et al., 2014; Prideaux and Warburton, 2010; Szalay, 1994). Conversely, characteristic features of more complex and heterogenous habitats include the increased transverse surface area of the talotibial articulation, more oblique orientation of the continuous lower ankle joint, a reduced area of talofibula contact, a broad sustentaculum tali and smoothing of the ventromedial and dorsolateral facets of the calcaneocuboid articulation (Bishop, 1997; Flannery and Szalay, 1982; Janis et al., 2014; Warburton and Prideaux, 2010).

Visualisation of three-dimensional shape differences between extant large macropods in our sample demonstrates that *Osphranter* exemplify the characteristics of open adapted ankle bones while *Notamacropus* and *Wallabia* ankle bones and, to a lesser extent *Macropus*, exhibit the features of a more mobile ankle joint. These characteristics broadly align with the typical habitat preferences of these genera. Although subtle compared to the differences between more distantly related macropodids, our results indicate that similar ecomorphological adaptations described by Prideaux and Warburton (2010) can also be observed between extant large macropod ankle bones. Our research therefore suggests that large macropod ankle bones, particularly the calcaneus, have potential to be used as ecomorphological proxies for local habitat structure and patch choice by Indigenous hunters.

The taxonomic and allometric differences in large macropod ankle shape identified in our study largely relate to differences in the orientation, relative positioning and curvature of articular surfaces that are difficult to adequately describe with linear measurements. In our previous study using linear measurements, we proposed that an elongated astragalar neck and calcaneocuboid step could be an ontogenetic trait in macropods (Mein et al., 2022). However, visualisation of three-dimensional allometric patterns reveals that it is the increasing anterior curvature of the medial trochlear crest that reduces this measurement in larger astragali rather than substantial differences to the anterior projection of the astragalar head. On the calcaneus, differences in the angle between the dorsolateral and dorsomedial calcaneocuboid facets increases the length of this linear measurement in smaller specimens. This highlights how piloting a GMM-based analysis, such as this study, may prove useful to ‘tease out’ the characteristics that distinguish groups and support the design of more effective linear morphometric protocols (see Viacava et al., 2023).

Although we have demonstrated that shape differences do occur between the four macropod genera in our sample, differences described between some taxa are subtle. Such subtle diagnostic attributes may be difficult to use in analyses that rely on visual comparison alone and may be easily confused with allometric variation. In these cases, machine learning can be used to correctly classify large macropod ankle bones to genus with a very high degree of accuracy and is an effective tool for the identification of morphologically similar specimens. The non-linear methods evaluated in our study consistently outperformed LDA by producing more balanced within-group sensitivity scores and were less likely to misclassify *Notamacropus* specimens. Non-linear statistical classifiers are better suited for use with high-dimensional GMM data (Dryden and Mardia, 1993), and make fewer assumptions about underlying data structure. This can be particularly beneficial to real world applications where sample sizes are often small and uneven (Cole et al., 2022; Courtenay et al., 2019; Moclán et al., 2023). Non-linear machine learning models which permit the inclusion of a high number of predictors can be particularly useful where shape differences between groups of interest are subtle or require a landmark approach to effectively describe. In these cases, discriminating shape information may be contained on low-ranked principal components that can be difficult to include in traditional discriminant function analyses when within-group sample sizes are low. Tolerance of research aims to within-group misclassification should drive model choice and different models should be tested on new data to evaluate differences in within-group misclassification (Soda et al., 2017).

Overall, statistical classification of macropod ankle bones via machine learning is a highly promising use case for palaeozoological applications. A larger training sample may permit classification of macropod specimens to species, or alternatively, classification to habitat type without the need for taxonomic identification. Future research should investigate the potential application of these methods to other elements of the macropod postcranial skeleton.

Statistical classification of palaeozoological specimens using morphometric predictors can also help to address many of the issues regarding quality assurance and data replicability that have long concerned zooarchaeologists (see Driver, 1992; Lyman, 2019; Rea, 1986; Wolverton, 2013). It is widely understood that identifying palaeozoological specimens by visually comparing their anatomical attributes with comparative reference collections, even by experienced analysts, includes a degree of uncertainty and error (Gobalet, 2001). Yet inter-observer error, or an analyst’s uncertainty, can be difficult to quantify or report.

Machine learning using morphometric data provides one means of increasing the analytical transparency in zooarchaeological specimen identification. As we have demonstrated here, overall accuracy of each machine learning model can be tested, the probability of correct group classification, and the posterior probabilities of group membership for each unknown specimen can be efficiently reported and, importantly, critically evaluated. Specimen identifications are easily replicated between analysts and where additional comparative data becomes available, training samples can be augmented to improve model accuracy and generalisability (Kovarovic et al., 2011; Kuhn and Johnson, 2016). Machine learning is increasingly used in archaeology for taxonomic or ecomorphological classification of specimens, bone fracture patterns and bone surface modifications (see Cucchi et al., 2021; Domínguez-Rodrigo et al., 2020; Moclán et al., 2023). Although statistical classification is yet to be widely adopted in archaeology in Australia, our study shows that these tools have significant potential to help with the taxonomic identification of morphologically similar macropod postcranial specimens from palaeozoological contexts.

The application of GMM and machine learning to zooarchaeological problems of specimen identification is becoming increasingly accessible, as specimen digitisation and landmark data capture becomes more cost efficient. However, data sharing is essential, as the time costs associated with the collation of the necessary training samples is the primary barrier for the adoption of these methods. Therefore, the digitised models, landmark training data and an easily replicable R code used in our study are provided open access for future research applications (see data availability statement).

## Conclusion

In this paper, we have demonstrated how the application of GMM can improve our understanding of taxonomically diagnostic characteristics between the ankle bones of closely related large macropod taxa. The four genera examined in this study are geographically widespread and there is substantial scope for the diagnostic characteristics identified here to be applied to palaeozoological specimens from multiple regions in Australia. Although we did not attempt to characterise diagnostic attributes beyond the genus level, in some cases species level identifications using these attributes could be well justified based on careful examination of specimen size and species biogeography. Combined with machine learning, these postcranial remains that have previously been considered unreliable to identify beyond the family level using visual comparative methods, can be statistically classified to genus with a high degree of accuracy. Macropods are one of the most speciose marsupial groups in Australia and the development of tools that allow us to differentiate their postcranial remains beyond broad family and size classes would create opportunities to better evaluate hypotheses regarding diet breadth, foraging efficiency and resource depression in Australia. Future research should investigate the impact specimen fragmentation has on classification accuracy to expand the applicability of these methods to palaeozoological problems. The combination of GMM and machine learning provides a powerful, quantitative, and replicable method for the identification of morphologically similar taxa and has high potential to be successfully applied to other postcranial elements and other speciose marsupial groups in Australia.

## Acknowledgements

Our thanks go to the collections managers who facilitated access to modern specimens used as the training sample: Heather Janetzki and Will Goulding (QM); David Stemmer (SAM); Kenny Travouillon (WAM); Gavin Dally and Barry Russell (MAGNT); Sandy Ingleby and Harry Parnaby (AM); and Leo Joseph and Chris Wilson (ANWC). Thanks also to the Ozboneviz project and Jacob van Zoelen for the production and use of several three-dimensional models used in the training sample.

## Data Availability

All landmark training data and R code to replicate this study can be found at www.github/ErinMein/Sorting-the-Mob. Digitised 3D models of all specimens in the training samples can be found at www.morphosource.org/projects/000524598.

## References

1. Adams, D.C., Collyer, M.L., Kaliontzopoulou, A., Baken, E., 2022. Geometric Morphometric Analyses of 2D/3D Landmark Data. https://github.com/geomorphR/geomorph.

2. Adams, D.C., Otarola-Castillo, E., 2013. geomorph: an R package for the collection and analysis of geometric morphometric shape data. Methods in Ecology and Evolution 4, 393–399. 10.1111/2041-210X.12035.

3. Adams, D.C., Rohlf, F.J., Slice, D.E., 2013. A field comes of age: geometric morphometrics in the 21st century. Hystrix 24(1), 7–14. 10.4404/hystrix-24.1-6283.

4. Aplin, K., Manne, T., Attenbrow, V., 2016. Using a 3-stage burning categorization to assess post-depositional degradation of archaeofaunal assemblages: Some observations based on multiple prehistoric sites in Australasia. Journal of Archaeological Science: Reports 7, 700–714. 10.1016/j.jasrep.2015.11.029.

5. Arriaza, M.C., Domínguez-Rodrigo, M., 2016. When felids and hominins ruled at Olduvai Gorge: A machine learning analysis of the skeletal profiles of the non-anthropogenic Bed I sites. Quaternary Science Reviews 139, 43–52. 10.1016/j.quascirev.2016.03.005.

6. Barnett, C.H., 1970. Talocalcaneal movements in mammals. Journal of Zoology 160(1), 1–7. 10.1111/j.1469-7998.1970.tb02893.x

7. Baudinette, R.V., 1977. Locomotory Energetics in a Marsupial, *Setonix brachyurus*. Australian Journal of Zoology 25, 423–428.

8. Baylac, M., Frieß, M., 2005. Fourier descriptors, Procrustes superimposition, and data dimensionality: An example of cranial human populations. In Slice, D.E.. (ed.), Modern Morphometrics in Physical Anthropology, pp. 145–165. Kluwer Academic/Plenum Publishers, New York.

9. Bennett, M. B., 1987. Fast locomotion of some kangaroos. Journal of Zoology, 212(3), 457–464. 10.1111/j.1469-7998.1987.tb02916.x

10. Bird, D.W., Bliege Bird, R., Codding, B.F., 2009. In Pursuit of Mobile Prey: Martu Hunting Strategies and Archaeofaunal Interpretation. Antiquity 74(1), 3–29. http://www.jstor.com/stable/25470536

11. Bird, D.W., Codding, B., Bliege Bird, R., 2013. Megafauna in a continent of small game: Archaeological implications of Martu Camel hunting in Australia’s Western Desert. Quaternary International 297, 155–166. 10.1016/j.quaint.2013.01.011

12. Bishop, N., 1997. Functional Anatomy of the Macropodid Pes. Proceedings of the Linnean Society of New South Wales 117, 17–50.

13. Bochenski, Z.M., 2008. Identification of skeletal remains of closely related species: the pitfalls and solutions. Journal of Archaeological Science 35(5), 1247–1250. 10.1016/j.jas.2007.08.013.

14. Bookstein, F.L., 1991. Morphometric tools for landmark data: geometry and biology. Cambridge University Press, Cambridge.

15. Bookstein, F.L., 1989. Principal Warps: Thin-Plate Splines and the Decomposition of Deformations. IEEE Transactions on Pattern Analysis Machine Intelligence 11(6), 567–585. 10.1109/34.24792.

16. Broughton, J.M., Cannon, M.D., Bartelink, E.J., 2010. Evolutionary Ecology, Resource Depression, and Niche Construction Theory: Applications to Central California Hunter-Gatherers and Mimbres-Mogollon Agriculturalists. Journal of Archaeological Method and Theory 17, 371–421. 10.1007/s10816-010-9095-7

17. Broughton, J.M., Cannon, M.D., Bayham, F.E., Byers, D.A., 2011. Prey Body Size and Ranking in Zooarchaeology: Theory, Empirical Evidence, and Applications from the Northern Great Basin. American Antiquity 76, 403–428. 10.7183/0002-7316.76.3.403.

18. Broughton, J.M., Broughton, M.J., Cole, K.E., Dalmas, D.M., Brenner Coltrain, J., 2023. Late Holocene tule elk (*Cervus canadensis nannodes*) resource depression and distant patch use in central California: Faunal and isotopic evidence from King Brown and the Emeryville Shellmound. Journal of Anthropological Archaeology 71, 101512. 10.1016/j.jaa.2023.101512

19. Cardini, A., Polly, P.D., 2020. Cross-validated Between Group PCA Scatterplots: A Solution to Spurious Group Separation? Evolutionary Biology 47, 85–95. 10.1007/s11692-020-09494-x.

20. Cignoni, P., Callieri, M., Corsini, M., Dellepiane, M., Ganovelli, F., Ranzuglia, G., 2008. MeshLab: an Open-Source Mesh Processing Tool. In Sixth Eurographics Italian Chapter Conference. pp. 129–136.

21. Codding, B.F., 2011. “Any Kangaroo?” on the Ecology, Ethnography and Archaeology of Foraging in Australia’s Arid West. Unpublished PhD thesis, Stanford University.

22. Cole, K.E., Yaworsky, P.M., Hart, I.A., 2022. Evaluating statistical models for establishing morphometric taxonomic identifications and a new approach using Random Forest. Journal of Archaeological Science 143, 105610. 10.1016/j.jas.2022.105610

23. Collyer, M.L., Sekora, D.J., Adams, D.C., 2015. A method for analysis of phenotypic change for phenotypes described by high-dimensional data. Heredity 115, 357–365. 10.1038/hdy.2014.75

24. Courtenay, L.A., 2023. Can We Restore Balance to Geometric Morphometrics? A Theoretical Evaluation of how Sample Imbalance Conditions Ordination and Classification. Evolutionary Biology 50, 90–110. 10.1007/s11692-022-09590-0

25. Courtenay, L.A., Yravedra, J., Huguet, R., Aramendi, J., Maté-González, M.Á., González-Aguilera, D., Carmen Arriaza, M., 2019. Combining machine learning algorithms and geometric morphometrics: A study of carnivore tooth marks. Palaeogeography, Palaeoclimatology, Palaeoecology 522, 28–39. 10.1016/j.palaeo.2019.03.007

26. Cucchi, T., Domont, A., Harbers, H., Leduc, C., Guidez, A., Bridault, A., Hongo, H., Price, M., Peters, J., Briois, F., Guilaine, J., Vigne, J.-D.D., 2021. Bones geometric morphometrics illustrate 10th millennium cal. BP domestication of autochthonous Cypriot wild boar (Sus scrofa circeus nov. ssp). Scientific Reports 11, 11435. 10.1038/s41598-021-90933-w

27. Dawson, T.J., 2012. Kangaroos, 2nd ed. CSIRO Publishing.

28. Dawson, T.J., 1995. Kangaroos: biology of the largest marsupials. University of New South Wales Press Ltd, Sydney.

29. Dawson, T. J., & Taylor, C. R., 1973. Energetic cost of Locomotion in Kangaroos. Nature, 246, 313–314. 10.1038/246313a0

30. Domínguez-Rodrigo, M., Cifuentes-Alcobendas, G., Jiménez-García, B., Abellán, N., Pizarro-Monzo, M., Organista, E., Baquedano, E., 2020. Artificial intelligence provides greater accuracy in the classification of modern and ancient bone surface modifications. Scientific Reports 10, 18862. 10.1038/s41598-020-75994-7

31. Drake, A.G., Klingenberg, C.P., 2008. The pace of morphological change: historical transformation of skull shape in St Bernard dogs. Proceedings of the Royal Society B: Biological Sciences. 275, 71–76. 10.1098/rspb.2007.1169.

32. Driver, J.C., 1992. Identification, Classification and Zooarchaeology. Circaea 9(1), 35–47.

33. Dryden, I.L., Mardia, K. V., 1993. Multivariate Shape Analysis. Sankhyā Indian Journal of Statistics Series A 55(3), 460–480. https://www.jstor.org/stable/25050954

34. Evin, A., Cucchi, T., Cardini, A., Strand Vidarsdottir, U., Larson, G., Dobney, K., 2013. The long and winding road: identifying pig domestication through molar size and shape. Journal of Archaeological Science 40, 735–743. 10.1016/j.jas.2012.08.005

35. Flannery, T.F., 1982. Hindlimb Structure and Evolution in the Kangaroos (Marsupialia: Macropodoidea), In Rich, P. V., Thompson, E.M., (eds.), The Vertebrate Fossil Record of Australasia, pp. 508–524. Monash University Offset Printing Unit, Melbourne.

36. Flannery, T.F., Szalay, F.S., 1982. Bohra paulae, a new giant fossil tree kangaroo (Marsupialia: Macropodidae) from New South Wales, Australia. Australian Mammalogy 5(2), 83–94. 10.1071/am82010.

37. Freedman, C. R., Rothschild, D., Groves, C., & Newman, A. E. M., 2020. *Osphranter rufus* (Diprotodontia: Macropodidae). Mammalian Species, 52(998), 143–164. 10.1093/mspecies/seaa011.

38. Garvey, J.M., 2010. Economic anatomy of the Bennett’s wallaby (Macropus rufogriseus): Implications for understanding human hunting strategies in late Pleistocene Tasmania. Quaternary International 211, 144–156. 10.1016/j.quaint.2009.07.006.

39. Gobalet, K.W., 2001. A Critique of Faunal Analysis; Inconsistency among Experts in Blind Tests. Journal of Archaeological Science 28(4), 377–386. 10.1006/jasc.2000.0564.

40. Goodall, C., 1991. Procrustes Methods in the Statistical Analysis of Shape. Journal of the Royal Statistical Society. Series B 53, 285–339. 10.1111/j.2517-6161.1991.tb01825.x.

41. Gould, R.A., 1969. Subsistence behaviour among the Western Desert Aborigines of Australia. Oceania 39(4), 253–274.

42. Guillerme, T., Weisbecker, V., 2019. landvR: Tools for measuring landmark position variation. 10.5281/zenodo.2620785.

43. Hair, J.F., Black, W.C., Babin, B.J., Anderson, R.E., 2018. Multivariate Data Analysis, 8th ed. Cengage.

44. Hanot, P., Guintard, C., Lepetz, S., Cornette, R., 2017. Identifying domestic horses, donkeys and hybrids from archaeological deposits: A 3D morphological investigation on skeletons. Journal of Archaeological Science 78, 88–98.

45. Hopwood, P. R., Butterfield, R. M., 1990. The locomotor apparatus of the crus and pes of the eastern grey kangaroo, *Macropus giganteus*. Australian Journal of Zoology, 38, 397–413. 10.1071/zo9900397.

46. Janis, C.M., Buttrill, K., Figueirido, B., 2014. Locomotion in Extinct Giant Kangaroos: Were Sthenurines Hop-Less Monsters? PLoS ONE 9(10), e109888. 10.1371/journal.pone.0109888.

47. Klingenberg, C.P., 1996. Multivariate Allometry. In Marcus, L.F., Corti, M., Loy, A., Naylor, G.J., Slice, D.E. (eds.), Advances in Morphometrics, pp. 23–49. Springer Science+Business Media, New York. 10.1007/978-1-4757-9083-2_3.

48. Klingenberg, C.P., 2016. Size, shape, and form: concepts of allometry in geometric morphometrics. Development Genes and Evolution 226, 113–137. 10.1007/s00427-016-0539-2.

49. Klingenberg, C.P., Monteiro, L.R., 2005. Distances and Directions in Multidimensional Shape Spaces: Implications for Morphometric Applications. Systematic Biology 54(4), 678–688. 10.1080/10635150590947258.

50. Kovarovic, K., Aiello, L.C., Cardini, A., Lockwood, C.A., 2011. Discriminant function analyses in archaeology: are classification rates too good to be true ? Journal of Archaeological Science 38, 3006–3018. 10.1016/j.jas.2011.06.028

51. Kuhn, M., 2022. Classification and Regression Training. https://github.com/topepo/caret/.

52. Kuhn, M., Johnson, K., 2016. Applied Predictive Modeling. Springer. 10.1007/978-1-4614-6849-3.

53. Lubinski, P.M., Lyman, R.L., Johnson, M.P., 2020. Blind Testing of Faunal Identification Protocols: A Case Study with North American Artiodactyl Stylohyoids. American Antiquity 85(4), 781–794. 10.1017/aaq.2020.45.

54. Lyman, R.L., 1992. Anatomical considerations of utility curves in zooarchaeology. Journal of Archaeological Science 19, 7–22. 10.1016/0305-4403(92)90003-L.

55. Lyman, R.L., 1994. Vertebrate Taphonomy. Cambridge University Press, Cambridge.

56. Lyman, R.L., 2019. Assumptions and Protocol of the Taxonomic Identification of Faunal Remains in Zooarchaeology: a North American Perspective. Journal of Archaeological Method and Theory 26, 1376–1438. 10.1007/s10816-019-09414-0.

57. Medit, 2019. ezScan 2017.

58. Mein, E. & Manne, T., 2021. Identifying marsupials from Australian archaeological sites: current methodological challenges and opportunities in zooarchaeological practice. Archaeology in Oceania 56, 133–141. 10.1002/arco.5234.

59. Mein, E., Manne, T., Veth, P., Weisbecker, V., 2022. Morphometric classification of kangaroo bones reveals paleoecological change in northwest Australia during the terminal Pleistocene. Scientific Reports 12, 18245. 10.1038/s41598-022-21021-w

60. Mitchell, D.R., Cairns, S.C., Koertner, G., Bradshaw, C.J.A., Saltré, F., Weisbecker, V., 2023. Differential developmental rates and demographics in red kangaroo populations separated by the dingo barrier fence. *Journal of Mammalogy*, gyad053. 10.1093/jmammal/gyad053

61. Mitteroecker, P., Bookstein, F., 2011. Linear Discrimination, Ordination, and the Visualization of Selection Gradients in Modern Morphometrics. Evolutionary Biology 38, 100–114. 10.1007/s11692-011-9109-8.

62. Moclán, A., Domínguez-García, Á.C., Stoetzel, E., Cucchi, T., Sevilla, P., Laplana, C., 2023. Machine Learning interspecific identification of mouse first lower molars (genus Mus Linnaeus, 1758) and application to fossil remains from the Estrecho Cave (Spain). Quaternary Science Reviews 299, 107877. 10.1016/j.quascirev.2022.107877

63. Morin, E., Ready, E., Boileau, A, Beauval, C., Coumont, M.P., 2017. Problems of Identification and Quantification in Archaeozoological Analysis, Part I: Insights from a Blind Test. Journal of Archaeological Method and Theory 24, 886–937. 10.1007/s10816-016-9300-4.

64. Morin, E., Meier, J., El Guennouni, K., Moigne, A.M., Lebreton, L., Rusch, L., Valensi, P., Conolly, J., Cochard, D., 2019. New evidence of broader diets for archaic *Homo* populations in the northwestern Mediterranean. Science Advances 5, eaav9106.

65. Nagaoka, L., 2019. Human Behavioral Ecology and Zooarchaeology. In: Prentiss, A.M. (ed.), Handbook of Evolutionary Research in Archaeology, pp. 231–253. Springer Nature, Cham. 10.1007/978-3-030-11117-5_12.

66. O’connell, J.F., Allen, J., 2012. The Restaurant at the End of the Universe: Modelling the colonisation of Sahul. Australian Archaeology 74, 5–17.

67. O’connell, J.F., Marshall, B., 1989. Analysis of kangaroo body part transport among the Alyawara of Central Australia. Journal of Archaeological Science 16(4), 393–405. 10.1016/0305-4403(89)90014-9

68. Peters, C., Richter, K.K., Manne, T., Dortch, J., Paterson, A., Travouillon, K.J., Louys, J., Price, G.J., Petraglia, M., Crowther, A., Boivin, N., 2021. Species identification of Australian marsupials using collagen fingerprinting. Royal Society Open Science 8, 211229. 10.1098/rsos.211229.

69. Peters, C., Wang, Y., Vakil, V., Cramb, J., Dortch, J., Hocknull, S., Lawrence, R., Manne, T., Monks, C., Rössner, G. E., Ryan, H., Siversson, M., Ziegler, T., Louys, J., Price, G. J., Boivin, N., & Collins, M. J., 2023. Bone collagen from subtropical Australia is preserved for more than 50,000 years. Communications Earth and Environment, 4(1), 1–8. 10.1038/s43247-023-01114-8

70. Press, A.J., 1989. The abundance and distribution of Black Wallaroos Macropus bernardus and Common Wallaroos Macropus robustus on the Arnhem Land Plateau Australia. In Grigg, G., Jarman, P., Hume, I., (eds.), Kangaroos, Wallabies and Rat-Kangaroos, pp. 783-786. Surrey Beatty & Sons Pty Limited, Chipping Norton.

71. Prideaux, G.J., Warburton, N.M., 2010. An osteology-based appraisal of the phylogeny and evolution of kangaroos and wallabies (Macropodidae: Marsupialia). Zoological Journal of the Linnean Society 159(4), 954–987. 10.1111/j.1096-3642.2009.00607.x.

72. R Development Core Team, 2023. R: A language and environment for statistical computing.

73. Rea, A.M., 1986. Verification and reverification: problems in archaeofaunal studies. Journal of Ethnobiology 6(1), 9–18.

74. Ritchie, E., 2007. The ecology and conservation of the antilopine wallaroo (Macropus antilopinus). Unpublished PhD thesis, James Cook University.

75. Rohlf, F.J., Slice, D.E., 1990. Extensions of the Procrustes Method for the Optimal Superimposition of Landmarks. Systematic Zoology 39, 40–59. 10.2307/2992207.

76. Salifu, D., Ibrahim, E.A., Tonnang, H.E.Z., 2022. Leveraging machine learning tools and algorithms for analysis of fruit fly morphometrics. Scientific Reports 12, 7208. 10.1038/s41598-022-11258-w.

77. Sanchez, P. M., 1974. The unequal group size problem in discriminant analysis. Journal of the Academy of Marketing Science 2, 629–633. 10.1007/BF02729456.

78. Schlager, S., Jefferis, G., Dryden, I.L., 2021. Calculations and Visualisations Related to Geometric Morphometrics. https://github.com/zarquon42b/Morpho.

79. Shining 3D, 2022. EinscanPro v2.6.0.9. [Software].

80. Soda, K.J., Slice, D.E., Naylor, G.J.P., 2017. Artificial neural networks and geometric morphometric methods as a means for classification: A case-study using teeth from *Carcharhinus* sp. (Carcharhinidae). Journal of Morphology 278, 131–141. 10.1002/jmor.20626.

81. Stratovan Corporation, 2018. *Stratovan Checkpoint* [Software].

82. Strauss, R.E., 2010. Discriminating groups of organisms. In Elewa, A.M.T. (ed.), Morphometrics for Nonmorphometricians, pp. 73–91. Springer, Berlin. 10.1007/978-3-540-95853-6_4

83. Stiner, M.C., Munro, N.D., Surovell, T., 2000. The Tortoise and the Hare: Small-Game Use, the Broad-Spectrum Revolution, and Paleolithic Demography. Current Anthropology 41(1), 39–79.

84. Szalay, F.S., 1994. Evolutionary history of the marsupials and an analysis of osteological characters. Cambridge University Press, Cambridge.

85. Van Dyck, S., Strahan, R., (eds.) 2008. The Mammals of Australia. 3rd edition. Sydney, Reed New Holland.

86. Viacava, P., Blomberg, S., Weisbecker, V., 2023. Geometric morphometrics out-perform linear-based methods in the taxonomic resolution of a mammalian species complex. Ecology and Evolution 13, e9698. 10.1002/ece3.9698

87. Warburton, N.M., Prideaux, G.J., 2010. Functional pedal morphology of the extinct tree-kangaroo Bohra (Diprotodontia: Macropodidae). In Coulson, G., Eldridge, M., (eds.), Macropods: The Biology of Kangaroos, Wallabies, and Rat-Kangaroos. pp. 137–151. CSIRO Publishing, Collingwood.

88. Weisbecker, V., Beck, R.M.D., Guillerme, T., Harrington, A., Lange-Hodgson, L., Lee, M.S.Y., Mardon, K., Phillips, M., 2023. Multiple modes of inference reveal less phylogenetic signal in marsupial basicranial shape compared to the rest of the cranium. Philosophical Transactions of the Royal Society B 378(1880). 10.1098/rstb.2022.0085.

89. Wolfhagen, J., Price, M.D., 2017. A probabilistic model for distinguishing between sheep and goat postcranial remains. Journal of Archaeological Science: Reports 12, 625– 631. 10.1016/j.jasrep.2017.02.022.

90. Wolverton, S., 2013. Data Quality in Zooarchaeological Faunal Identification. Journal of Archaeological Method and Theory 20(3), 381–396. 10.1007/s10816-012-9161-4.

91. Zeder, M.A., Lapham, H.A., 2010. Assessing the reliability of criteria used to identify postcranial bones in sheep, *Ovis*, and goats, *Capra*. Journal of Archaeological Science 37(11), 2887–2905. 10.1016/j.jas.2010.06.032.

