## Supplementary for "Sorting the mob: using geometric morphometrics and machine learning to differentiate kangaroo postcrania for zooarchaeological applications"

### Supplementary Information

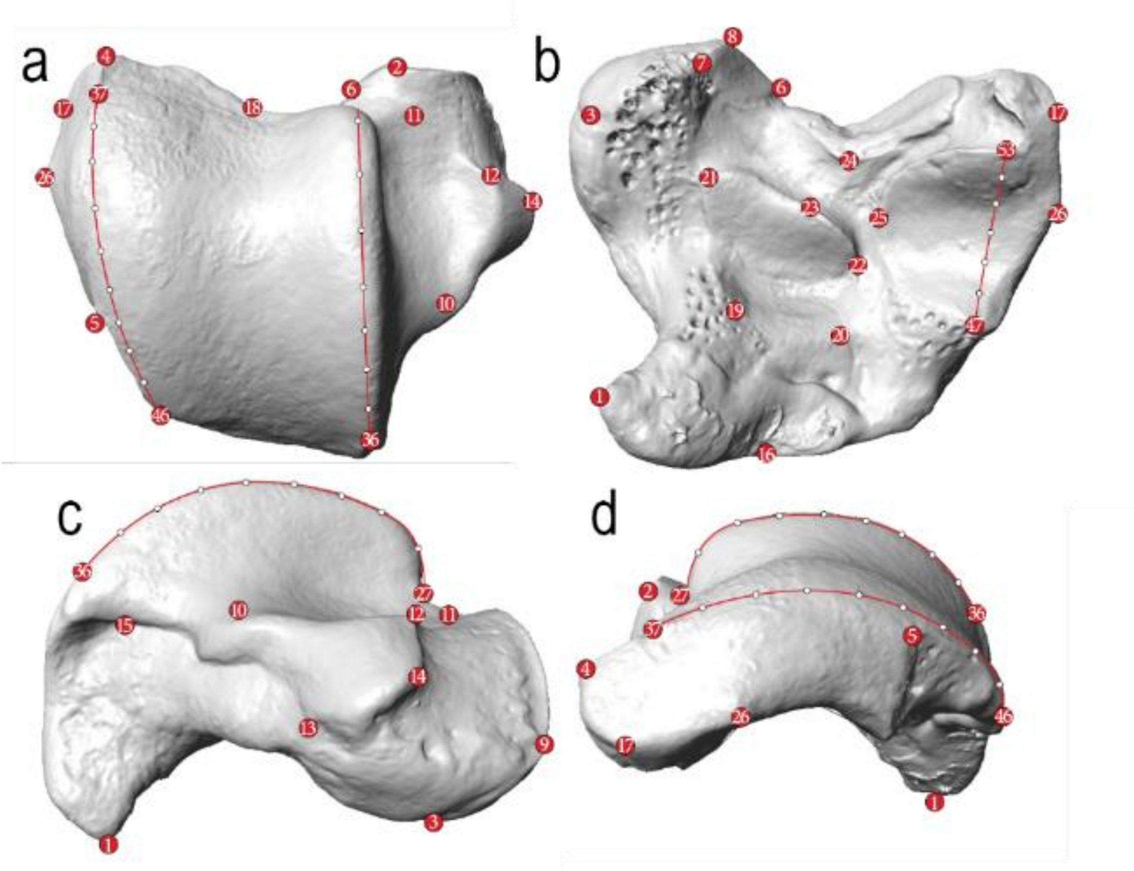


**Figure S1** Large macropod astragalus landmarking protocol. **a)** superior view, **b)** inferior view, **c)** medial view and **d)** lateral view. Red circles indicate fixed landmarks, white circles indicate sliding semi-landmarks.

**Table S1** Definition of homologous landmarks on large macropod astragali. Note: anatomical directions and descriptions of ligament attachments for the definition of the landmarks follows Bishop (1997).

| Landmark | Definition |
|---|---|
| 1 | Most plantar point on the medial plantar tuberosity |
| 2 | Most antero-dorsal point on the talocuboid articulation on the lateral aspect of the astragalar head |
| 3 | Most plantar point on the talonavicular articulation |
| 4 | Most anterior point of the lateral trochlear crest |
| 5 | Most dorsal point along the posterior margin of the area of attachment for the lateral meniscus |
| 6 | Most dorso-lateral point of the talocuboid articulation along the junction with the adjoining talocuboid articular facet on the lateral aspect of the astragalar head |
| 7 | Most plantar point of the secondary talocalcaneal articulation on the lateral aspect of the astragalar head |
| 8 | Most antero-plantar point of on the talocuboid articulation along the junction with the adjoining talocuboid articular facet on the lateral aspect of the astragalar head |
| 9 | Point of maximum curvature along the medial margin of the talo-navicular facet |
| 10 | Point of maximum curvature along the posterior margin of the malleolar fossa |
| 11 | Point of maximum curvature along the anterior margin of the malleolar fossa |
| 12 | Most dorsal point of the junction between the medial process and the malleolar fossa |
| 13 | Most planto-posterior point of the medial process |
| 14 | Point of maximum curvature of the medial process |
| 15 | Point of maximum curvature of the dorsal margin of the rugose area of attachment for the posterior tibiotalar ligament |
| 16 | Junction of the posterior margin of the talotibial trochlea and medial plantar tuberosity |
| 17 | Most antero-lateral point of the lateral talocalcaneal articular facet |
| 18 | Point of maximum curvature of the anterior margin of the talotibial trochlea |
| 19 | Most medio-posterior point of the medial talocalcaneal articular facet |
| 20 | Most latero-posterior point of the medial talocalcaneal articular facet |
| 21 | Most medio-anterior point of the medial talocalcaneal articular facet |
| 22 | Lateral origin of the anterior margin of the medial talocalcaneal articular facet |
| 23 | Point of maximum curvature along the anterior margin of the medial talocalcaneal articular facet |
| 24 | Point of maximum curvature of the XXX facet/attachment?? |
| 25 | Point of maximum curvature of the medial margin of the lateral talocalcaneal articular facet |
| 26 | Most lateral point of the lateral talocalcaneal articular facet |
| 27 | The point of maximum concavity at the anterior point of the medial trochlear crest |
| 28-35 | Series of eight sliding semi-landmarks place at equidistant intervals along the apex of the medial trochlear crest between fixed landmarks 27 and 36. |
| 36 | Most posterior point of the medial trochlear crest at junction with the medial plantar tuberosity |
| 37 | The point of maximum concavity at the anterior point of the lateral trochlear crest |
| 38-45 | Series of eight sliding semi-landmarks place at equidistant intervals along the apex of the lateral trochlear crest between fixed landmarks 37 and 46. |
| 46 | Most posterior point of the lateral trochlear crest |

| | |
|---|---|
| 47 | Most posterior-plantar point of the lateral talocalcaneal articular facet |
| 48-52 | Series of five sliding semi-landmarks place at equidistant intervals along the curvature of the lateral talocalcaneal articular facet between fixed landmarks 47 and 53. |
| 53 | Most antero-plantar point of the lateral talocalcaneal articular facet |

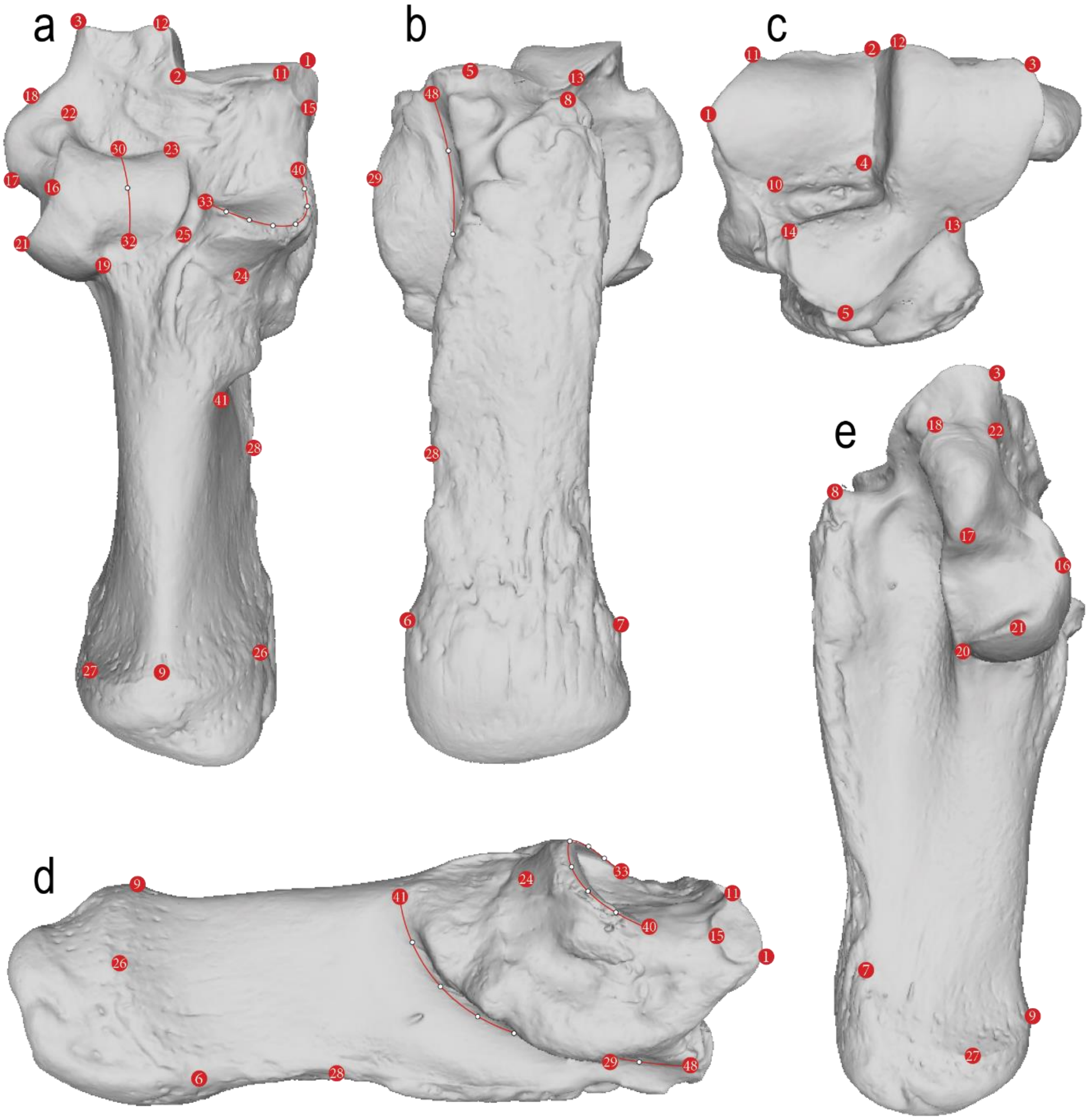


**Figure S2** Large macropod calcaneus landmarking protocol. **a)** superior view, **b)** inferior view, **c)** anterior view, **d)**, medial view and **e)** lateral view. Red circles indicate fixed landmarks, white circles indicate sliding semi-landmarks.

**Table S2** Definition of homologous landmarks on large macropod calcanea. Note: anatomical directions and descriptions of ligament attachments for the definition of the landmarks follows Bishop (1997).

| Landmark | Definition |
|---|---|
| 1 | Most medial point of the dorsomedial calcaneocuboid articular facet |

| | |
|---|---|
| 2 | Most dorsal point at the junction between the dorsolateral and dorsomedial articular facet |
| 3 | Dorsolateral corner of the dorsolateral articular facet |
| 4 | Most plantar point of the junction between the dorsolateral and dorsomedial articular facet |
| 5 | Most plantar point of the ventromedial calcaneocuboid articular facet |
| 6 | Medial posterior point of the rugose plantar surface along the margin of the epiphyseal suture |
| 7 | Lateral posterior point of the rugose plantar surface along the margin of the epiphyseal suture |
| 8 | Most antero-lateral point of the rugose plantar surface along the margin of the sulcus between the plantar surface and calcaneocuboid articulation |
| 9 | Most dorsal point of the epiphyseal suture on the tuber calcanei |
| 10 | Most plantar point of the dorsomedial calcaneocuboid articular facet |
| 11 | Dorsomedial corner of the dorsomedial calcaneocuboid articular facet |
| 12 | Dorsomedial corner of the dorsolateral calcaneocuboid articular facet |
| 13 | Plantar-lateral corner of the dorsolateral calcaneocuboid articular facet at the point of maximum concavity between the dorsolateral and ventromedial facets |
| 14 | Dorsomedial corner of the ventromedial calcaneocuboid articular facet |
| 15 | Point of maximum convexity of the secondary calcaneotalar facet on the medial surface of the dorsomedial calcaneocuboid articular facet |
| 16 | Most dorsal point of the area of attachment for the posterior calcaneofibular ligament where it adjoins the lateral margin of the lateral calcaneotalar articular facet |
| 17 | Most posterior point of the process for the attachment of the anterior calcaneofibular ligament along the margin with the area of attachment for the posterior calcaneofibular ligament |
| 18 | Most anterior point of the process for the attachment of the anterior calcaneofibular ligament |
| 19 | Junction of the latero-posterior corner of the lateral calcaneotalar articular facet and area of attachment for the posterior talocalcaneal ligament |
| 20 | Most plantar point of the calcaneofibular articular facet |
| 21 | Most lateral point of the calcaneofibular articular facet |
| 22 | Point of maximum concavity along the lateral margin of the fossa anterior to the lateral calcaneotalar articular facet |
| 23 | Medio-anterior corner of the lateral calcaneotalar articular facet at the junction with the fossa |
| 24 | Most posterior point of the posterior side of the medial calcaneotalar articular facet |
| 25 | Most medio-anterior point of the attachment area for the posterior talocalcaneal ligament in the sulcus between the medial and lateral calcaneotalar articular facets |
| 26 | Point of maximum convexity along the medial margin of the epiphyseal suture on the tuber calcanei |
| 27 | Point of maximum convexity along the lateral margin of the epiphyseal suture on the tuber calcanei |
| 28 | Point of maximum concavity along the medial margin of the rugose plantar surface on the tuber calcanei |
| 29 | Most plantar point on the sustentaculum tali |
| 30 | The mid-point of the anterior margin of the convex surface of the lateral calcaneotalar articular facet |
| 31 | A sliding semi-landmark placed equidistant between fixed landmarks 30 and 32. |

| | |
|---|---|
| 32 | The mid-point of the posterior margin of the convex surface of the lateral calcaneotalar articular facet |
| 33 | The most lateral point along the crest of the medial calcaneotalar articular facet |
| 34-39 | Series of six sliding semi-landmarks place at equidistant intervals along the crest of the medial calcaneotalar articular facet between fixed landmarks 33 and 40. |
| 40 | Antero-medial origin of the crest of the medial calcaneotalar articular facet |
| 41 | Most dorsoposterior point of the junction of the sustentaculum tali and tuber calcanei |
| 42-47 | Series of six sliding semi-landmarks place at equidistant intervals along the curved junction of the sustentaculum tali and tuber calcanei between fixed landmarks 41 and 48. |
| 48 | Most antero-plantar point of the junction of the sustentaculum tali and tuber calcanei |

**Table S3** Pairwise comparison of allometric slopes of large macropod astragali and calcanea.

| | d | UCL (95%) | Z | Pr > d |
|---|---|---|---|---|
| *Astragali* | | | | |
| *Macropus ~ Notamacropus* | 0.30074 | 0.334841 | 0.797614 | 0.206 |
| *Macropus ~ Osphranter* | 0.189246 | 0.201041 | 1.163822 | 0.123 |
| *Notamacropus ~ Osphranter* | 0.294723 | 0.29628 | 1.615101 | 0.057 |
| *Calcanea* | | | | |
| *Macropus ~ Notamacropus* | 0.246337 | 0.253908 | 1.501863 | 0.069 |
| *Macropus ~ Osphranter* | 0.203822 | 0.169086 | 2.901976 | 0.003 |
| *Notamacropus ~ Osphranter* | 0.232076 | 0.223518 | 1.949858 | 0.026 |

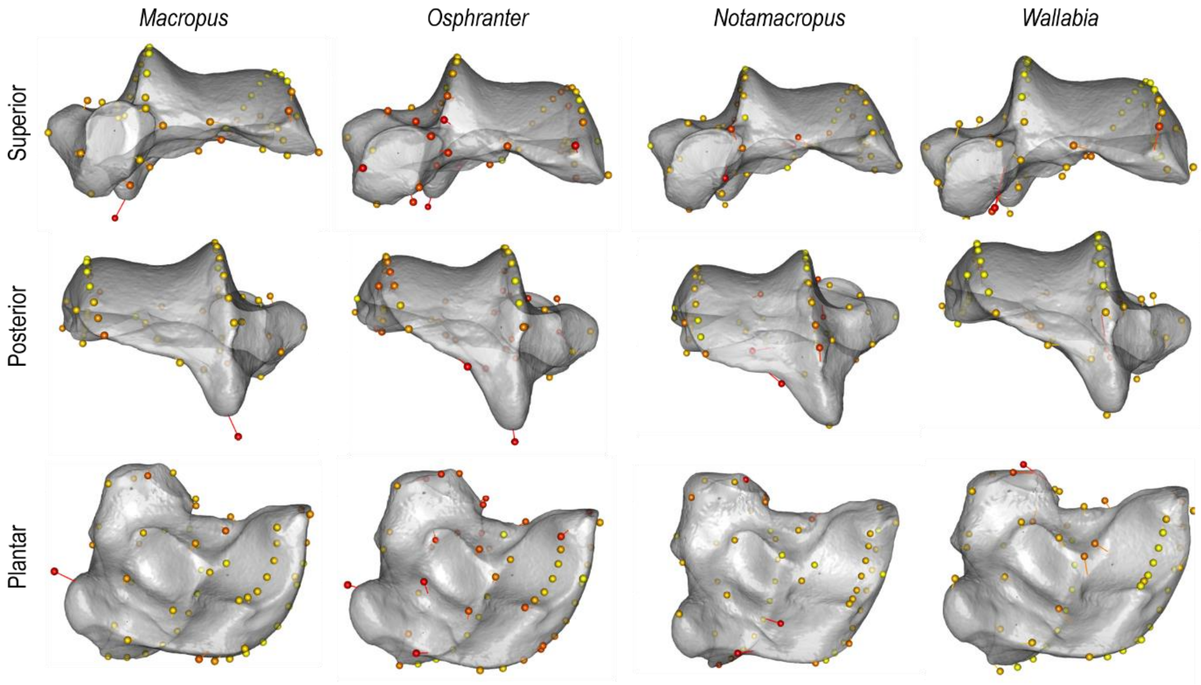


**Figure S3** Visualisation of allometric shape variation in the macropod astragalus by taxonomic group. Spheres are the position of one landmark in small specimens, lines represent the displacement of the same landmark in large specimens. Red spheres indicate areas of greatest variation and yellow spheres areas of least variation. Models are representation of small specimen shape warped along the predicted allometric trajectory for each genus.

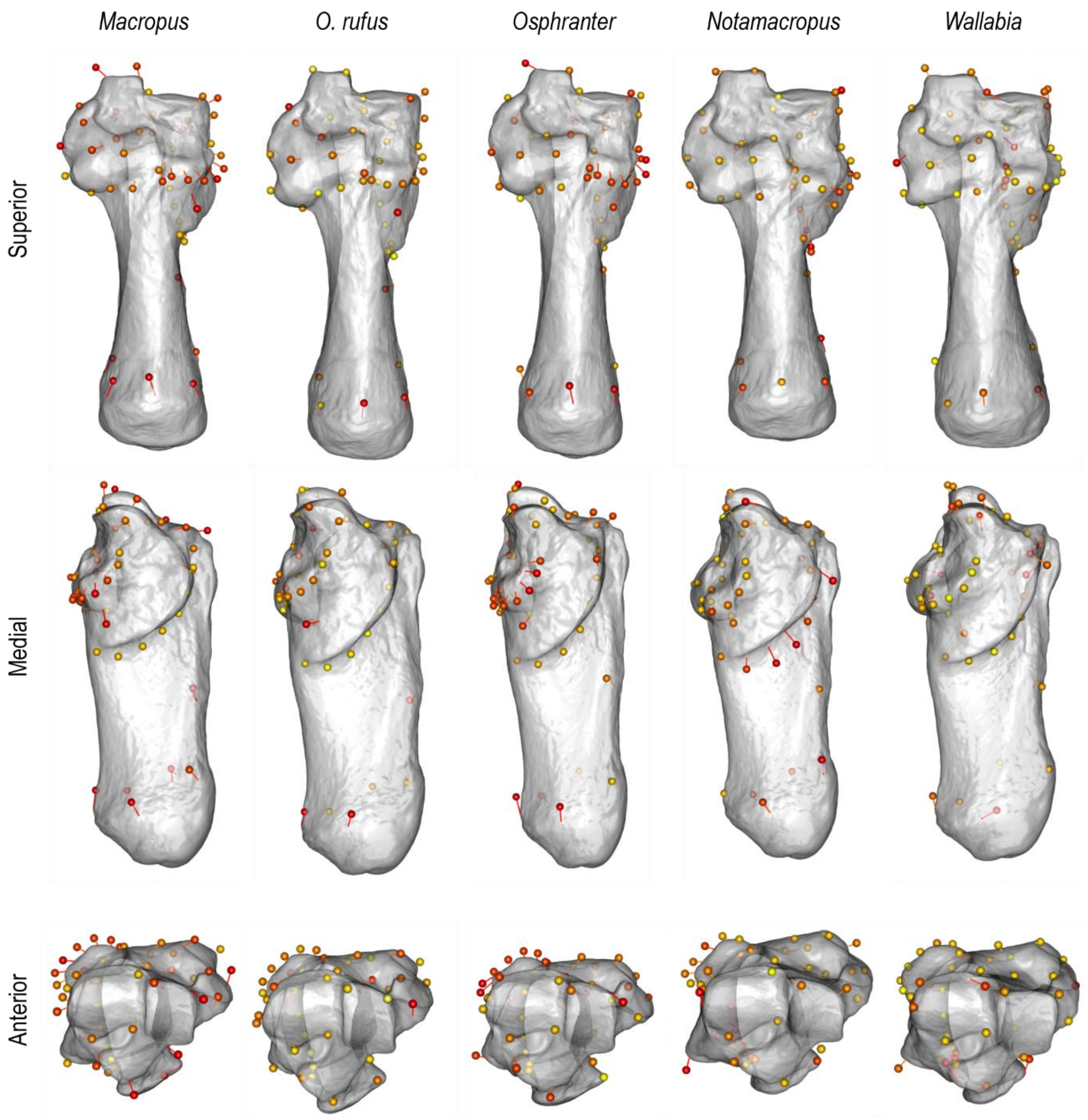


**Figure S4** Visualisation of allometric shape variation in the macropod calcaneus by taxonomic group. Spheres are the position of one landmark in small specimens, lines represent the displacement of the same landmark in large specimens. Red spheres indicate areas of greatest variation and yellow spheres areas of least variation. Models are representation of large specimen shape warped along the predicted allometric trajectory of each genus.

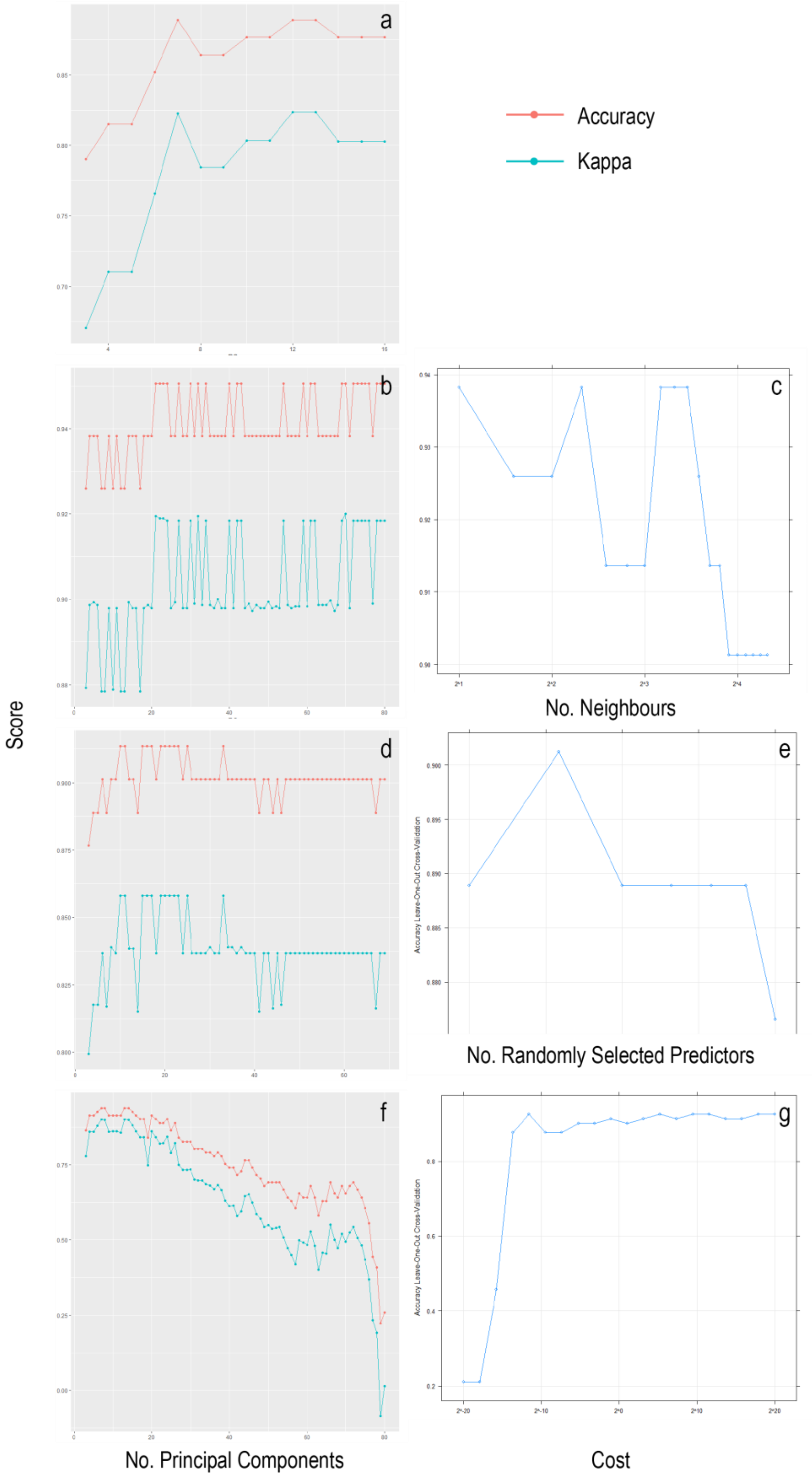


**Figure S5** Results of tuning the machine learning models to statistically classify macropod astragali. **Left)** Model accuracy and Kappa scores for astragalus classification against number of Principal Components (L), showing performance of each model with increasing PCs. **Right)** Accuracy of each model across the tuning parameters after number of Principal Components are optimised. **a)** LDA model; **b-c)** k-NN algorithm; **d-e)** RF model; and **f-g)** SVM model.

**Table S4** Cross validated confusion matrices for the statistical classification of large macropod astragali.

| | | Observed | | | |
|---|---|---|---|---|---|
| | | *Macropus* | *Notamacropus* | *Osphranter* | *Wallabia* |
| | *Linear Discriminant Analysis* | | | | |
| **Predicted** | *Macropus* | 17 | 0 | 0 | 0 |
| | *Notamacropus* | 0 | 9 | 0 | 3 |
| | *Osphranter* | 0 | 0 | 43 | 0 |
| | *Wallabia* | 0 | 8 | 1 | 0 |
| | *K Nearest Neighbour* | | | | |
| | *Macropus* | 17 | 1 | 0 | 0 |
| | *Notamacropus* | 0 | 14 | 0 | 3 |
| | *Osphranter* | 0 | 2 | 44 | 0 |
| | *Wallabia* | 0 | 0 | 0 | 0 |
| | *Random Forest* | | | | |
| | *Macropus* | 16 | 1 | 0 | 0 |
| | *Notamacropus* | 1 | 14 | 1 | 3 |
| | *Osphranter* | 0 | 2 | 43 | 0 |
| | *Wallabia* | 0 | 0 | 0 | 0 |
| | *Support Vector Machine* | | | | |
| | *Macropus* | 17 | 0 | 2 | 0 |
| | *Notamacropus* | 0 | 17 | 1 | 3 |
| | *Osphranter* | 0 | 0 | 41 | 0 |
| | *Wallabia* | 0 | 0 | 0 | 0 |

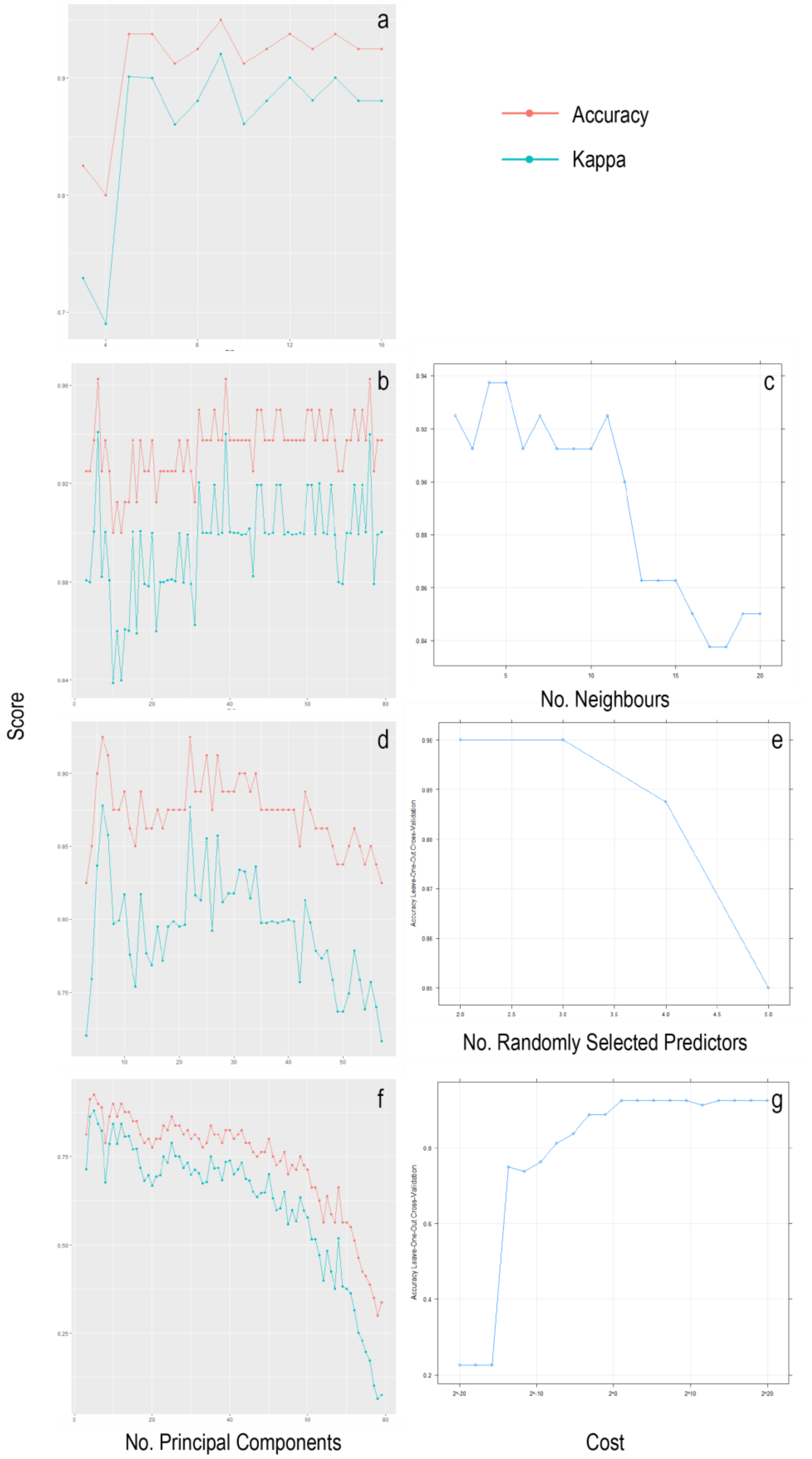


**Figure S6** Results of tuning the machine learning models to statistically classify macropod calcanea. **Left)** Model accuracy and Kappa scores for astragalus classification against number of Principal Components (L), showing performance of each model with increasing PCs. **Right)** Accuracy of each model across the tuning parameters after number of Principal Components are optimised. **a)** LDA model; **b-c)** k-NN algorithm; **d-e)** RF model; and **f-g)** SVM model.

**Table S5** Cross validated confusion matrices for the statistical classification of large macropod calcanea.

| Predicted | Observed: *Macropus* | *Notamacropus* | *Osphranter* | *Wallabia* |
|---|---|---|---|---|
| *Linear Discriminant Analysis* | | | | |
| *Macropus* | 18 | 2 | 0 | 0 |
| *Notamacropus* | 0 | 13 | 0 | 0 |
| *Osphranter* | 0 | 0 | 42 | 0 |
| *Wallabia* | 0 | 2 | 0 | 3 |
| *K Nearest Neighbour* | | | | |
| *Macropus* | 18 | 0 | 1 | 0 |
| *Notamacropus* | 0 | 16 | 1 | 3 |
| *Osphranter* | 0 | 0 | 40 | 0 |
| *Wallabia* | 0 | 1 | 0 | 0 |
| *Random Forest* | | | | |
| *Macropus* | 16 | 0 | 1 | 0 |
| *Notamacropus* | 0 | 16 | 0 | 3 |
| *Osphranter* | 2 | 0 | 41 | 0 |
| *Wallabia* | 0 | 1 | 0 | 0 |
| *Support Vector Machine* | | | | |
| *Macropus* | 18 | 1 | 1 | 0 |
| *Notamacropus* | 0 | 15 | 0 | 3 |
| *Osphranter* | 0 | 1 | 41 | 0 |
| *Wallabia* | 0 | 0 | 0 | 0 |

*Supplementary Files*

**Video S1** Visualisation of the differences in large macropod astragali between mean genus shapes. Reference mesh warped to mean group shape along PC1-PC3 of a cross validated bgPCA.

*Supplementary Files*

**Video S2** Visualisation of the differences between large macropod calcanea between mean genus shapes. Reference mesh warped to mean group shape along PC1-PC3 of a cross validated bgPCA.

*Supplementary Files*

**Video S3** Visualisation of allometric variation in astragalus shape between small and large specimens in each taxonomic group. Reference mesh warped between minimum and maximum shapes along 'PredLine' argument of function 'plotAllometry' in the package geomorph.

*Supplementary Files*

**Video S4** Visualisation of allometric variation in calcaneus shape between small and large specimens in each taxonomic group. Reference mesh warped between minimum and maximum shapes along 'PredLine' argument of function 'plotAllometry' in the package geomorph
